# Differential modulation of host cell Serine and Arginine-Rich (SR) protein and apoptotic factor BCL-2 gene expressions during Chandipura and Dengue virus infections

**DOI:** 10.64898/2026.11.14.766452

**Authors:** Dibyakanti Mandal

## Abstract

Regulation of alternative splicing (AS) is important in controlling cellular functions. More than 95% human genes are alternatively spliced and AS is regulated by close to 100 splicing factors. Line of evidences suggest that virus infections alter the AS pattern of numerous host genes. In this study we have investigated Chandipura virus (CHPV) induced alteration of host cell SR protein gene expressions and determined if these changes have any effect on apoptotic factor BCL-2 levels,. CHPV is a negative stranded RNA virus which causes fever, brain encephalitis and death among young children with a case-fatality rate of 50%. Few studies were done in past to understand the roles of host factors in CHPV pathogenesis and no study was done on virus induced changes in AS. Our study showed that there is a down-regulation of SR proteins SRp55 and SRp75 expression levels significantly decreased in the CHPV infected Vero E6 cells compared to uninfected cells. For a comparison, we also looked at the levels of SRp55 and SRp75 in dengue virus (DENV) infected cells. Our results showed that SRp75 levels is unchanged whereas SRp55 levels increased in DENV infected cells. We have also determined the levels of large and small splice variants of apoptotic gene BCL-2 to find out possible correlation between the altered splicing factor expressions and virus infected cell death via apoptosis. Our data indicates that in CHPV infected cells, BCL-2 large splice variant expressions decreased as compared to uninfected cells. This is the first study of CHPV mediated alteration of splicing factor expressions and these results warrant further study with other viruses to better understand the global changes in host splicing events, cell survival, host immune response during virus infection.

## Introduction

Virus infection leads to numerous changes in the transcriptome of host cell. Viruses try to gain access of the host cell machinery for productive infection and at the same time host cells make efforts to restrict the virus replication through differential expression of diverse sets of genes including genes of innate immunity (1–4). Viruses overcome these restrictions by inhibition of host gene expressions or modulation of expression levels. Dengue, HCV were known to inhibit innate immune response pathway by inhibiting interferon productions (5–7). Alternative splicing (AS) are important biological events in eukaryotic cells which regulate the gene expressions at post-transcription level (8–10). Alteration of AS leads to changes in cell function, signaling pathways, upregulation or downregulation of innate immune genes and cell survival (11–15). Recent study has shown that Dengue virus infection causes altered splicing pattern of SAT1 gene and further experiments indicated that the alteration is mediated through downregulation of cellular splicing factor RBM10 (16). AS is regulated by *cis*-acting elements present on the genes and splicing factors such as SR proteins, hnRNPs (17–22). Evidences suggest that viruses modulate the AS patterns of host genes, which has many consequences that decide the fate of both the virus and the host (23–25). Few studies were done to determine the changes in expression levels of splicing factors during virus infection, but not much known how these changes affect cellular AS pattern and key functions of the cells (26,27). Studies have shown that conversion of pro-apoptotic to apoptotic activity of some factors are due to production of splice variants through AS (28–31). As viruses are able to induce apoptosis of infected cells, it is important to know the mechanism of these pathways, whether modulation of AS by viruses has any role in conversion of apoptotic factors. In the present study we have investigated the expressions of SR protein genes in Chandipura virus infected cells. CHPV is a negative stranded RNA virus belonging to Rhabdovirus family. CHPV is an emerging pathogen of the Indian subcontinent and in some of the African countries (32,33). Since first reported in 1965 (34), several outbreaks were reported from the Indian states of Maharashtra, Gujrat, Bihar (35–38)). CHPV disseminates to the central nervous system via blood, causes fever, brain encephalitis among young children below 14 years of age and the case-fatality rate is more than 50% among the infected individuals. At present there is no targeted treatment for CHPV encephalitis, and clinical management is done by fluid and electrolyte balance, paracetamol, and diclofenac. Limited studies have been done till date to find out the host factors and their roles in CHPV pathogenesis (39,40). It was recently shown that CHPV infected neuronal cells undergoes rapid apoptosis and apoptotic factor FAS mediated signaling pathway is involved in the death of infected cells (41). It may be assumed that other apoptotic factors are also associated with the CHPV induced apoptosis. We hypothesize that CHPV infections to cells modulate the AS of one or more apoptotic genes to support productive infection. To test our hypothesis, we have studied the changes in expression levels of several SR protein genes and apoptotic factors BCL-2 in CHPV infected cells. These results were compared with Dengue virus, another RNA virus, to better understand the alteration pattern in viruses from different families.

## Materials and methods

### Cells and viruses

Vero E6 cells were used in all the experiments. Cells were maintained in DMEM media in presence of 10% FBS and 1x antibiotics (Penicillin-Streptomycin) mixture. Stock of CHPV Indian strain was used in all the experiments (Obtained from Dr. D.J. Chattopadhyay Laboratory, University of Kolkata). Titer of the virus stock was ∼10^8^/ml. Dengue virus infected cells in Trizol were obtained from Dr. S Samal, Translational Health Sciences and Technology Institute, Faridabad.

### CHPV infection

1 million Vero E6 cells were seeded in 6-well plates 24h before infection. Cells were infected at a m.o.i (multiplicity of infection) of 0.01 in serum free DMEM media for 2h at 37°C. Cells were washed with PBS twice and incubated at 37°C in complete DMEM media containing 10% FBS. Uninfected cells were used as control and were treated similarly except virus infection. 24h later infected and uninfected cells were scrapped and stored in -80 deg until used. . RNA was extracted from stored cells. **and cDNA <u>prepared.RNA extraction, cDNA preparation and PCR amplification of SR protein genes:</u>**

For RNA extraction cells were mixed with 0.5 ml Trizol (Invitrogen) and then RNA was extracted following manufacturer’s protocol. cDNAs from infected and uninfected cells was prepared from extracted RNA using cDNA synthesis kit (Applied Biosystems) following manufacturer’s protocol and used as template to amplify SR proteins genes using specific primers (Listed in table 1) (42). PCR amplification was performed at 95 deg 5 min, 35 cycles of 95 deg 15 sec, 55 deg 30 sec and 72 deg 1 min, followed by 72 deg 5 min. PCR amplified products were analyzed on 1.5% agarose gel.

**Table 1.**
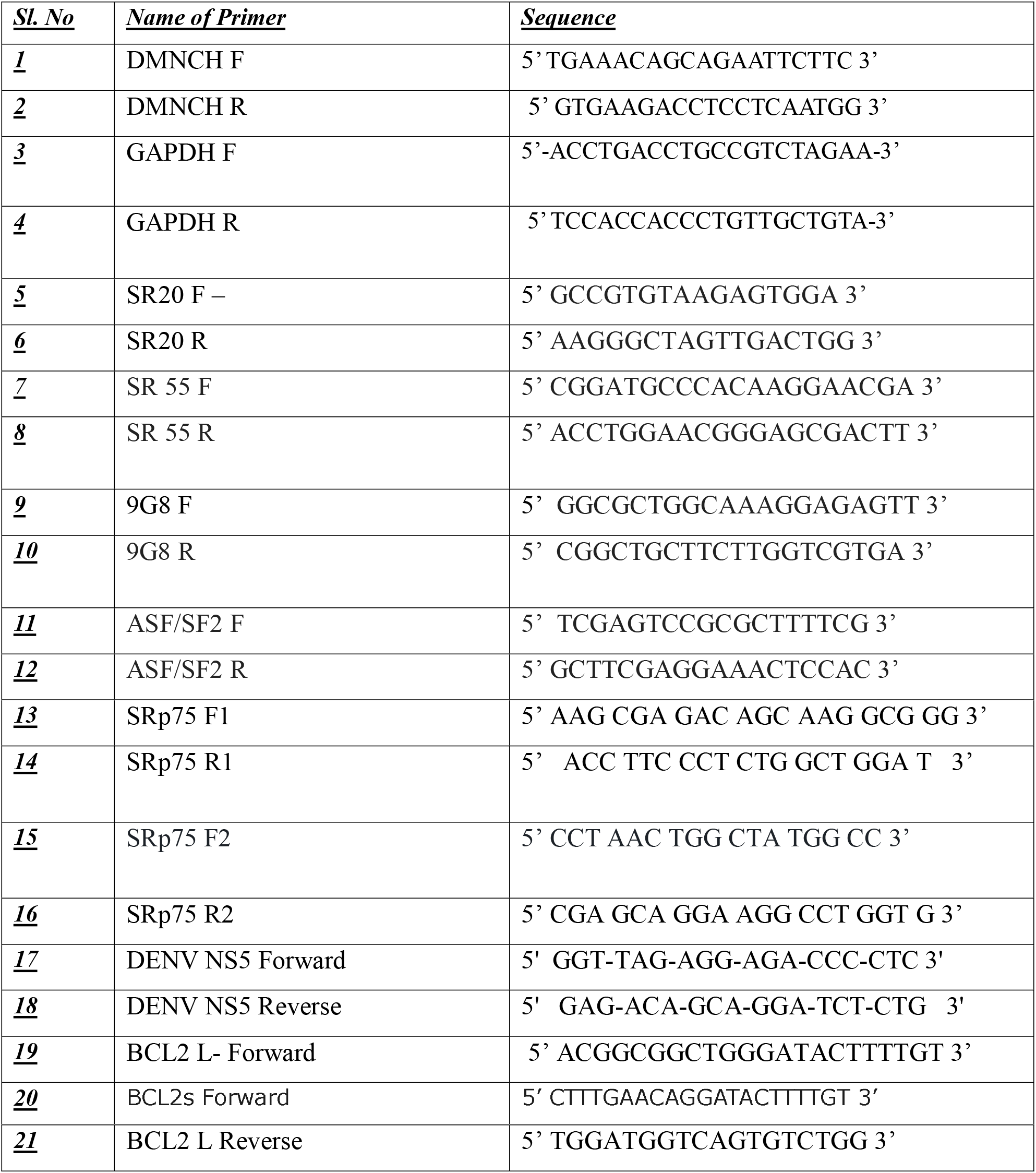

### Analysis of SR protein and BCL-2 genes by quantitative real-time PCR

For quantitative measurement of gene expression levels of SR proteins genes and also to confirm the PCR based results shown in Fig 1, we have performed a SYBR Green based real-time PCR assays. cDNA synthesized and used for PCR based assays were also used as template for quantitative PCR. Real-time PCR was done in presence of 1x SYBR Green PCR mix (Bio-Rad), 200 nM each of forward and reverse primers and 2.5 μl cDNA in a total volume of 20 μl. Reaction was performed at 25 deg 10 min, 1 cycle at 95 deg 2 min, 40 cycles of 95 deg 30 sec and 60 deg 1 min. We looked at the SRp55 and SRp75 genes as these gene showed noticeable changes in conventional PCR as shown in figure 1. We have also analysed the levels of BCL-2 large and small splice variants using targeted primers (Table 1). Ct values obtained were analyzed and comparison was made between infected and uninfected cells. GAPDH gene was used as internal control. Relative changes in expression of SR proteins genes were determined using ΔΔCt and 2^–ΔΔCt^ formula and compared (43) and plotted.

**Figure 1.**
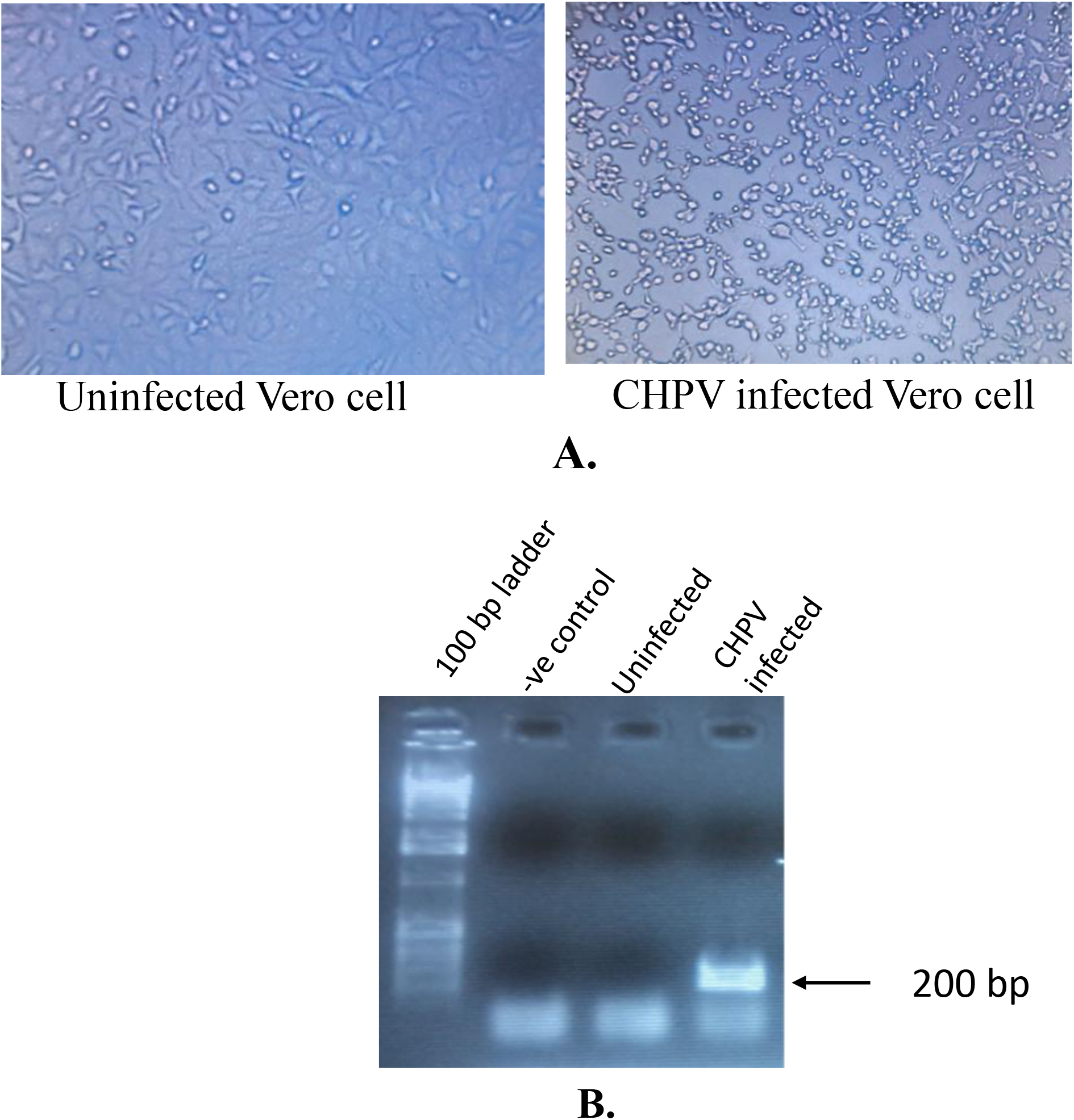
1a. CHPV infection in Vero E6 cells and observed cytopathic effects. Infected Vero E6 cells were observed under microscope (right) and compared with uninfected cells (left) 1b. cDNAs obtained from infected and uninfected cells were PCR amplified using CHPV N (nucleocapsid) specific primers and analyzed on 1.5% Agarose gel.

## Results

### CHPV infection induces altered splicing factor expression in vero cells

We have studied the expressions levels of cellular splicing factors, the SR protein genes in Chandipura virus infected Vero E6 cells and compared to that in uninfected controls. As mentioned in the materials method section, cDNAs prepared were used as template for PCR amplification of SRp55, 9G8, SRp20 and ASF/SF2 genes. We used GAPDH as internal control. To confirm that the Vero cells were infected with CHPV, 24 hr post-infection we looked at the cells under microscope. Clear cytopathic effects (cell rounding) were observed in CHPV infected cells, whereas no cytopathic effects were present in uninfected cells (Figure 1A). Infection was further confirmed by PCR amplification of CHPV N (nucleocapsid) gene using specific primers (table 1). As shown in figure 1A, analysis of PCR amplified products clearly showed a 200 bp N specific band in infected cell and no band present in PCR product in uninfected cells (Figure 1B).

Next, we compared the levels of SRp55, 9G8, SRp20, SRp75 and ASF/SF2 genes in CHPV infected cells to that in uninfected cells. We looked at the expressions of GAPDH in both infected and uninfected cells as an internal control. As shown in figure 2a, GAPDH levels are same in infected and uninfected cells (Fig. 2a lane 2 and 3). We found that SRp55 expressions were significantly affected in virus infected cells. SRp55 specific 265 bp band cells was absent infected cell but visible in uninfected cells, which indicates downregulation of SRp55 gene (Fig 2b, lane 5 and 6) in CHPV infection. Lanes 8 and 9 in figure 2a shows PCR amplified 9G8 gene products in uninfected and infected Vero cells, respectively. A decrease in level of 9G8 was noticed in infected cells. No major change in gene expression levels was noticed for SRp20 as shown in lane 11 and 12 in fig 2a. Small decrease in the levels of ASF/SF2 was noticed as shown in figure 2b (lane 2 -uninfected, lane 3-infected). We have used two different primer sets to look at the changes in the expression levels of SRp75 (Fig 2c). Result showed no PCR amplified SRp75 products in infected cells (Lanes 3 and 6), whereas a clear band is present in uninfected cells (lanes 2 and 5). We used ImageJ software (https://ij.imjoy.io/) to quantify the band intensity of PCR products. Figure 2c shows the densitometry comparison of DNA band intensities of PCR products obtained from uninfected and infected cells. The graph clearly shows noticeable decreases in SRp55 levels in infected cells as compared to uninfected cells, whereas the GAPDH (internal control) levels are same. Quantitative real-time PCR was done to confirm above results and also to determine relative amounts of expressed SR protein genes. We measured the quantity of SRp55 and SRp75, the genes which were found to be decreased in infected cells from the PCR and gel electrophoresis analysis. Real-time PCR analysis showed that expression levels of SRp55 andSRp75 decreased by 40% and 70% respectively in CHPV infected cells compared to uninfected cells (Fig. 2). These results are summarized in table 2.

**Table 2:** Summary of changes in expression levels of SRp55, SRp75 and BCL-2 in CHPV and DENV infected cells.

| **Splicing factor** | **CHPV infection** | ***p values*** | **DENV infection** | ***p value*** |
|---|---|---|---|---|
| SRp55 | 40% decrease | 0.017 | 40% increase | 0.0047 |
| SRp75 | 70% Decrease | 0.0119 | 10% increase | 0.568 |
| BCL2- L | 93% decrease | 0.003 | 86% decrease | 0.0083 |
| BCL-2 s | 72% decrease | 0.014 | 30% decrease | 0.05 |

### DENV infection does not alter SRp75 levels but increases SRp55 levels

We next examined the changes of splicing factor expression levels in dengue virus infected VeroE6 cells. Dengue virus infection was confirmed by PCR of infected cell cDNA using NS5 specific primers (sequences provided in table 1). PCR amplification of ∼200 pb product in DENV infected cells shown in figure 4A which confirmed that the cells are infected with the virus. We next looked at the SRp55, ASF/SF2 and SRp75 gene expression levels in DENV infected cells. PCR amplification and agarose gel electrophoresis showed that SRp75 and ASF/SF2 is expression levels were unchanged in DENV infection as compared to uninfected cells, whereas level of SRp55 was higher than uninfected cells (Figure 4A). Real-time PCR results also showed noticeable increase in the levels of SRp55 expressions in DENV infected cells as compared to uninfected and small increase in SRp75 expression levels (Fig 4B). Results are also summarized in table 2.

**Figure 2A.**
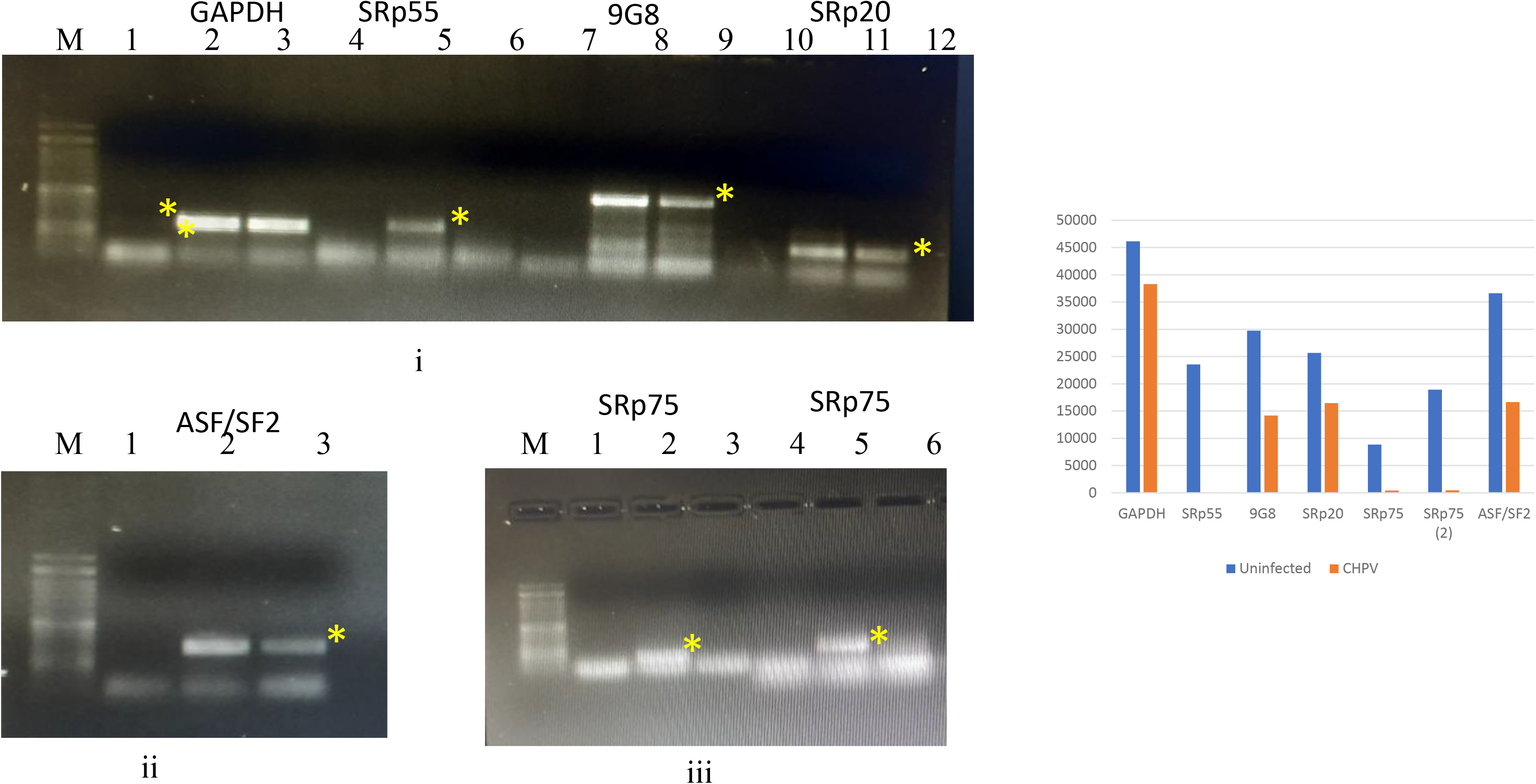
PCR amplification of SR protein genes from CHPV infected and uninfected Vero E6 cells. Figure 2A(i), 2A(ii). and 2A(iii). show agarose gel electrophoresis of PCR amplified products of different SR protein genes. Figure 2A(i), lane 2 and 3 – GAPDH uninfected and infected cell, respectively. Lane 5 and 6-SRp55 in uninfected and infected cells; lanes 8 and 9-9G8 from uninfected and infected cells; lanes 11 and 12 -SRp20 from uninfected and infected cells. In figure 2A(ii), lane 2 and 3 are ASF/SF2 in uninfected and infected cells. In figure 2A(iii), SRp75 amplification production. 2 and 5 are uninfected cells, 3 and 6 are from infected cells. Expected Product sizes are indicated by asterisks. Figure 2A(iv).-Densitometry analysis of bands in Fig 2a, 2b and 2c. Blue bars are showing band intensity from uninfected cells and yellow bars showing band intensity from infected cells,

**Figure 2B.**
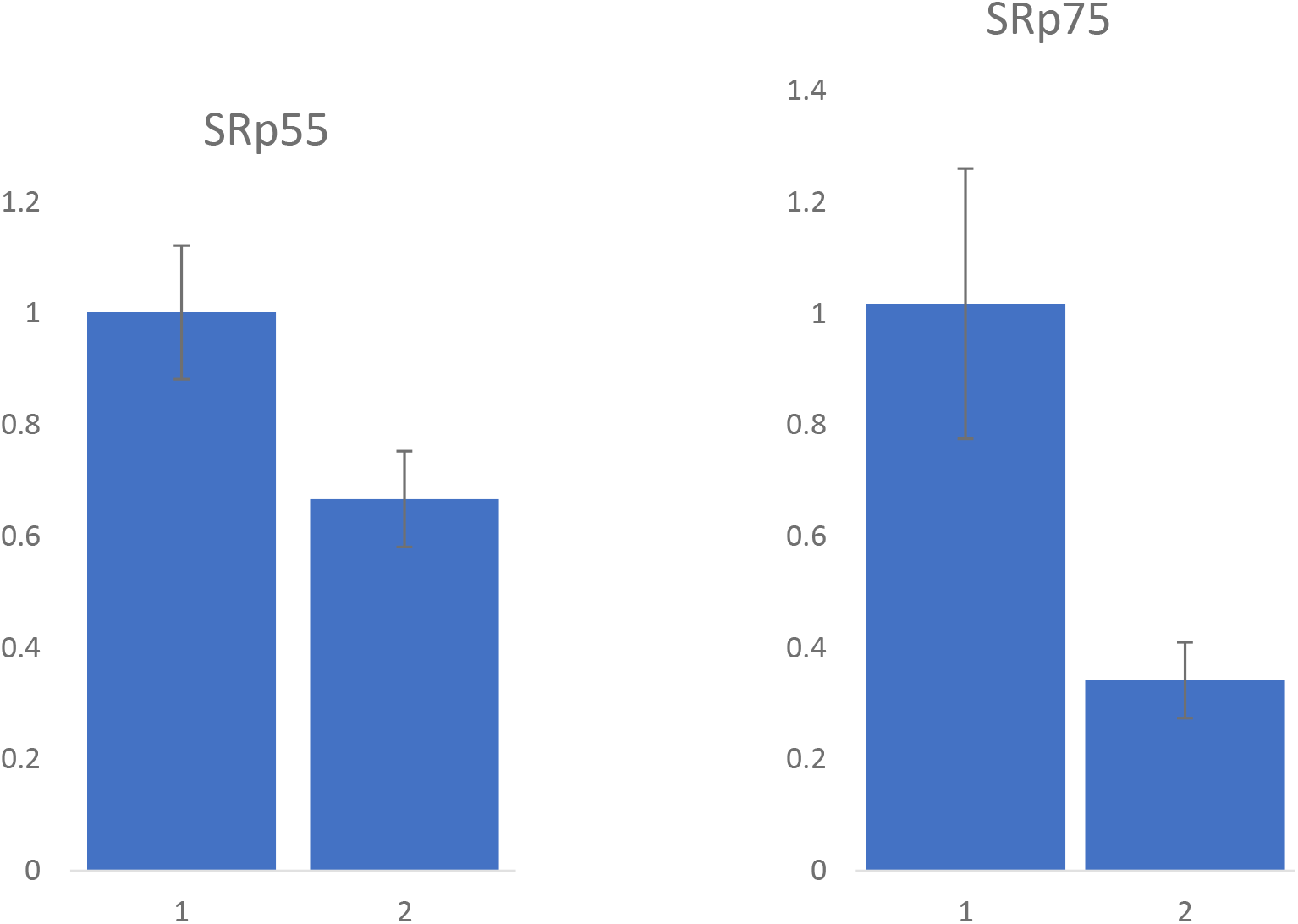
Quantitative real-time PCR of SRp55 and SRp75 genes from infected and uninfected cells. Relative amounts of the genes in infected and uninfected cells as compared to GAPDH were determined and plotted.

**Figure 3.**
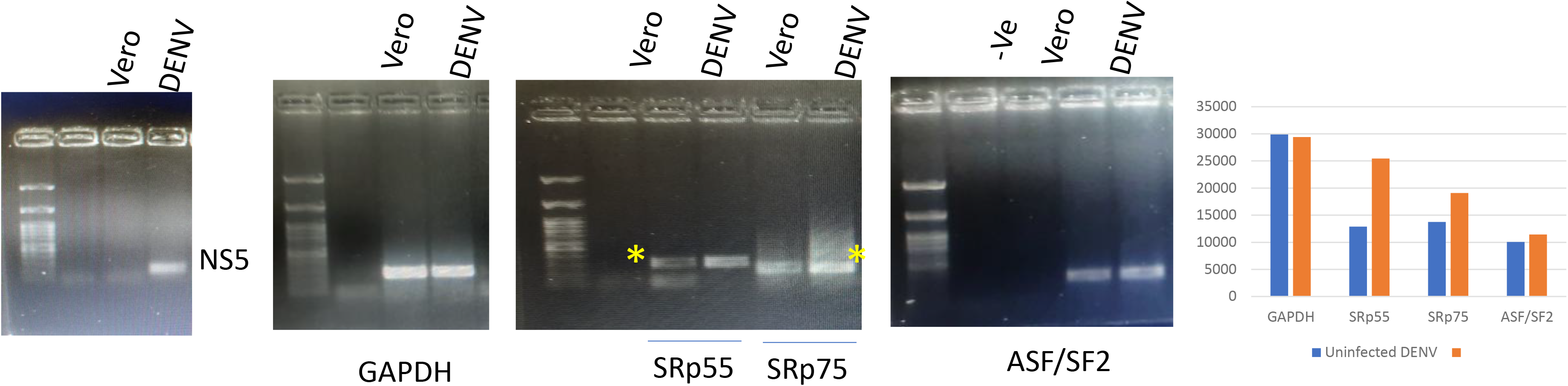
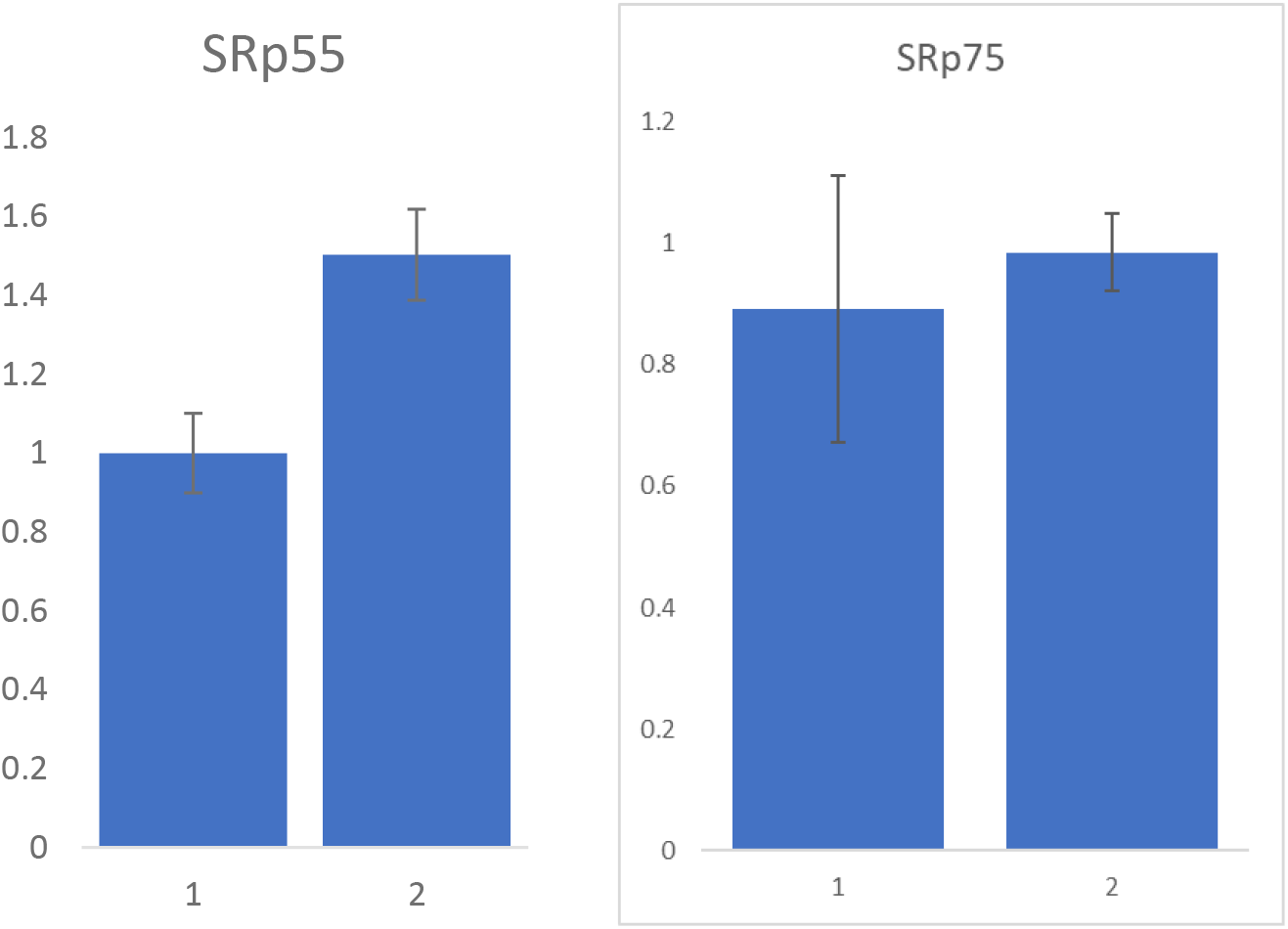
A. PCR amplification and agarose gel electrophoresis of DENV NS5, GAPDH, SRp55, SRp75 and ASF/SF2 genes from DENV infected and uninfected cells. B. Quantitative real-time PCR for SRp55 and SRp75 gene expressions in DENV infected and uninfected cells. Expected Product sizes are indicated by asterisks,

**Figure 4.**
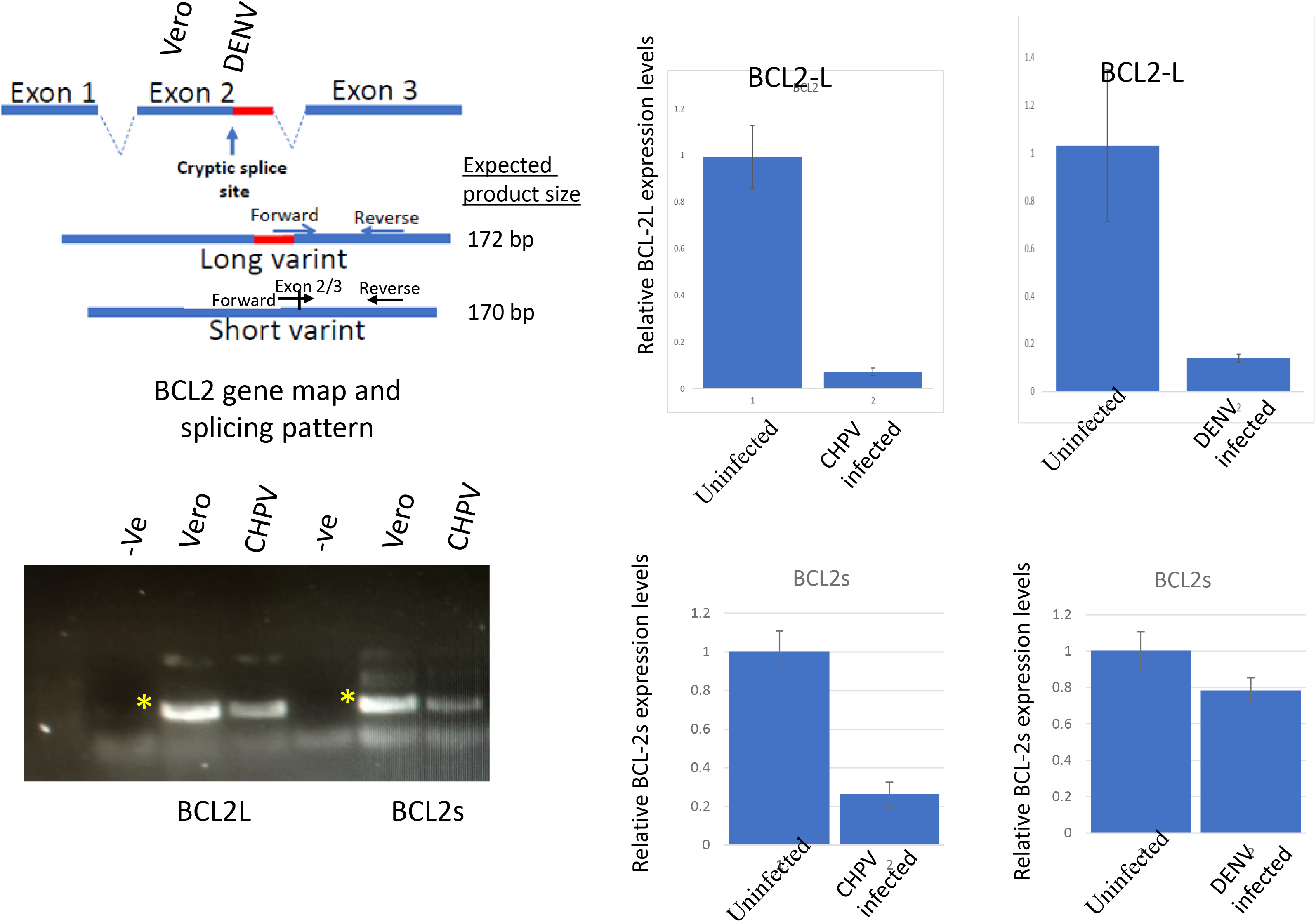
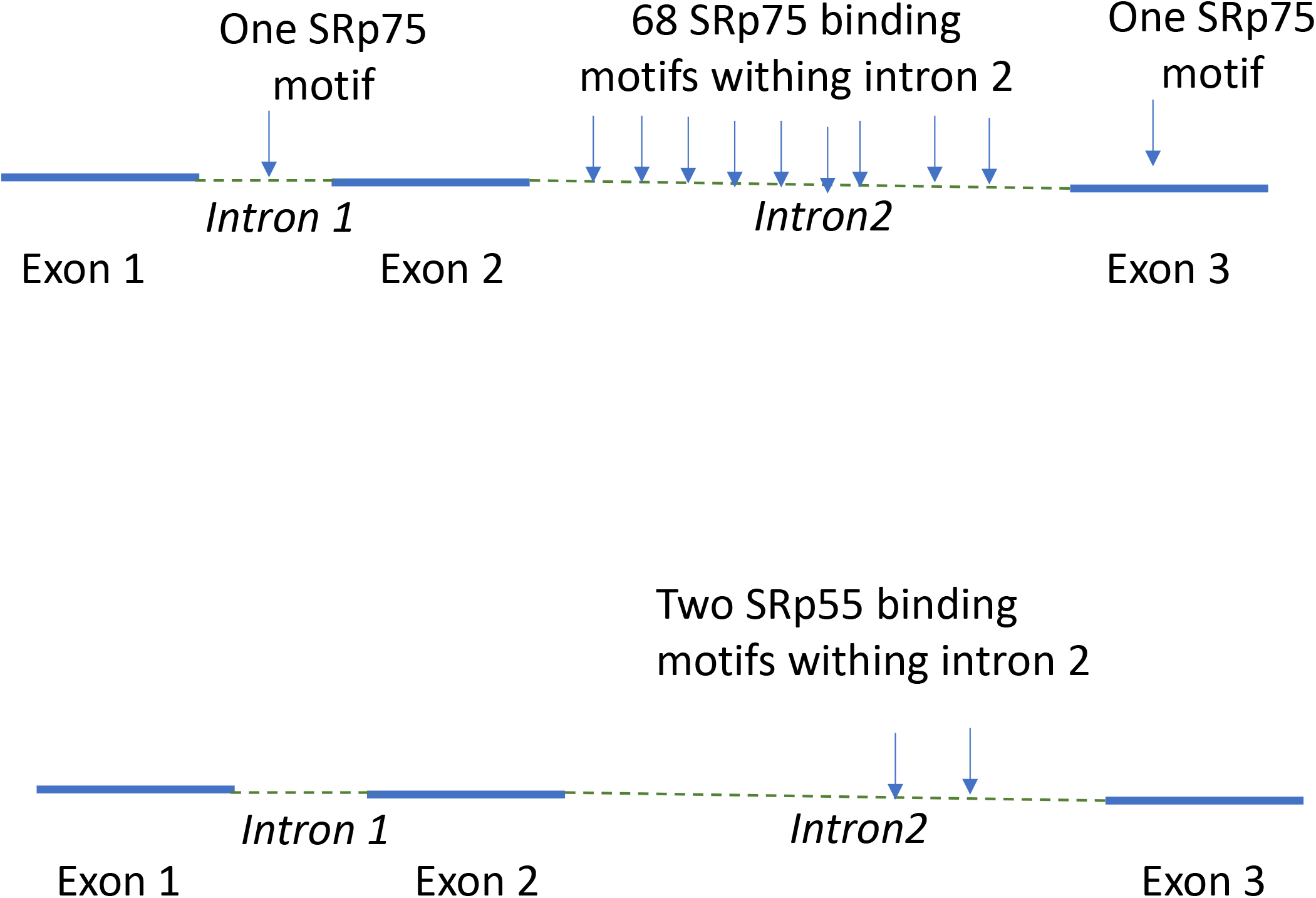
A. Apoptotic gene BCL-2 splicing pattern and position of primers used in amplification of BCL-2 large splicing variant (Left). Agarose gel pic showing PCR amplification of large and small variant of BCL2 gene from CHPV infected and uninfected cells. Expression levels of BCL-2 large splice variant and BCL-2 small splice variant in CHPV and DENV infected Vero cells compared to uninfected cells. (right, top and bottom respectively). **B.** SRp55 and SRp75 binding motif positions within BCL-2 gene.

### Decreased expressions of BCL-2 gene in CHPV infected cells

We were interested to know if altered expression pattern of SR protein genes has any correlation to cell apoptosis induced by CHPV and DENV. BCL-2 is an apoptotic factor which is expressed as a large and a small variant resulting from alternative splicing of the BCL-2 pre-mRNA. We have designed primer pairs, that are able to specifically amplify the large or small variant of the gene (Fig 5). Real-time PCR results showed that in both CHPV and DENV infected cells, levels of BCL-2 large variant were much lower compared to uninfected cells. In CHPV infected cells BCL-2 large variant is only 7% of the uninfected cell whereas in DENV infected cells the levels are 14% of uninfected cells (Fig 5). We have also looked at the expressions of small splice variant of BCL-2 or BCL2s. Our results showed that in CHPV infected cells there is a decrease in the levels of BCLs but proportion of small variant is 4-fold higher than large variant. Similar trend observed in Dengue virus infected cells where slight decrease in BCL-2s but expression is 5-fold higher compared to BCL-2 large variant (Figure 5B). GAPDH is the internal control and expression levels of both BCL-2 large and small variants are relative to GAPDH.

**Figure 5.**
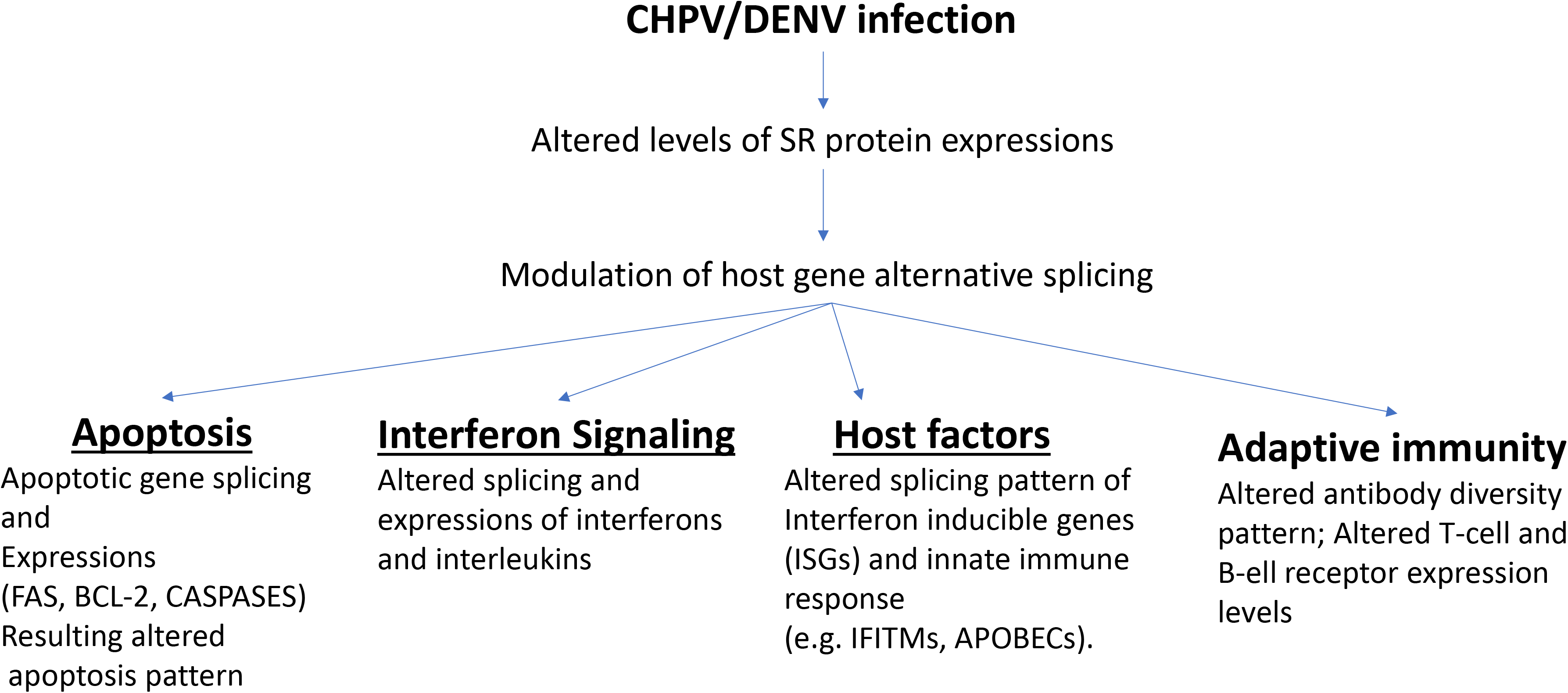
Predicted downstream effects of CHPV and DENV mediated changes in splicing factor expressions.

## Discussions

In the present study we have investigated the altered expressions of splicing factors in CHPV infected Vero E6 cells. We have looked at the expressions of SR proteins SRp55, 9G8, SRp20, ASF/SF2 and SRp75. We observed decreased levels of SRp55 and SRp75 in CHPV infected cells as compared to uninfected cells. Quantitative real-time PCR confirmed that SRp55 and SRp75 expression levels decreased by 40% and 70% respectively in the CHPV infected cells. CHPV is a negative stranded RNA virus and replication of the virus does not involve splicing of the virus gene. Therefore, our results suggest that virus induced modulation of SR protein genes has a deeper implication in virus pathogenesis. We have compared SRp55 and SRp75 expression levels in CHPV infected cells with DENV infection. In contrast to CHPV, in DENV infection SRp55 and SRp75 expression pattern is different. Small increase of SRp55 was observed in DENV infected cells and SRp75 level was unchanged. A previous study demonstrated that CHPV induced neuronal cell death takes place though FAS mediated signaling, one of the key apoptosis pathways (41). Another study showed that the overexpression of SRp55 changes the splicing pattern of FAS (44). SRp55 also regulates the pancreatic beta cell survival and lower expression of SRp55 leads to cell apoptosis (45). So, down-regulation of SRp55 and SRp75 in CHPV infected cell may have a correlation to infection induced cell death. We hypothesize that other apoptotic gene expressions and their altered splicing pattern is associated with CHPV mediated cell death and pathogenesis. Apoptotic factor BCL-2 is alternatively spliced and two major splice variants are detected in apoptotic cells. Our results showed that in CHPV infected cells large splice variant expression levels significantly decreased. Though BCL2s levels increased 4-fold compared to BCL-2 large variant, the levels decreased relative to GAPDH. This result suggesting that CHPV infection may have a dual effect on BCL-2 expressions; i) infection caused decrease in the expression levels of BCL-2 gene, and ii) it also alters the splicing pattern. Large splice variant of BCL-2 is anti-apoptotic and small variant is apoptotic (46). To find if there is any relationship between BCL-2 levels with decreased SRp55 and SRp75 in CHPV infected cells we have searched the binding motifs of these two SR proteins in BCL-2 genes. We have looked SRp55 motif ‘GCCGGA or TCCGGA’ and SRp75 binding motif ‘GAAAA or GAAGA’ (47,48). Our analysis shows that for SRp55, motif sequence GCCGGA is absent but two match for the sequence TCCGGA are present in BCL-2 gene (Fig 5B and Supplementary figure 1). For Srp75, 70 matches found for ‘GAAAA’ and 68 sequences of ‘GAAGA’ (Fig 5B and Supplementary figure 2. This indicates that downregulation of BCL-2 in CHPV infected cells is SRp75 dependent. In DENV infected cells also we observed decreased levels of BCL-2 large variant but decrease is lesser compared to CHPV. This study shows that CHPV and DENV have differential effects on splicing factor expression levels and also differently modulate the apoptotic factors. Further in-depth studies are needed to understand the complex interplay between cellular splicing factors and apoptotic gene-splicing pattern in virus infected cells. Not only apoptosis, there are other biological events which are dependent on alternative splicing of host genes (Illustrated in Figure 6). These include genes of interferons and ISGs (49). Interferon gene expressions are regulated through AS and alteration of splicing changes the expression levels of these factors, affecting innate immune response. ISGs, which are differently expressed and play roles in virus pathogenesis are also known to be alternatively spliced and little is known about their splice variants and antiviral activities. Natural splice variant of IFITM3 is known to determine the severity of Influenza infection and mortality. APOBEC, another well studied family of interferon induced host factor is alternatively spliced but detailed information on antiviral activities of their splice variants are limited. T-cell and B-cell mediated immunity provide targheted protection against viruses. Recent study results indicate that immune functions of T cell and B cells are dependent on the AS pattern of several genes in these cells which include T-cell and B-cell receptors, antibody variable region coding genes. This emerging information may help in understanding the acquired immune response during infection. In this aspect, our results may encourage others to conduct in-depth studies to know the splicing regulation of different levels of immune response against various pathogens. In conclusion, our findings may help to better understand the virus induced modulation of host genes’ alternative splicing and associated post-infection events within the cells. Novel anti-viral drugs may be designed to restore splicing events and normal cell function.

## Acknowledgement

Author thank Prof. D. J. Chattopadhayay, University of Calcutta, Kolkata for Chandipura Virus Indian strain and anti-N antibody. Author is thankful to Dr. S. Samal, THSTI, Faridabad and Dr. S Saraswat, Amity Institute of Virology for DENV infected cells.

## Conflict of interests

Author declare that there is no conflict of interest.

## Funding information

This work received no specific grant from any funding agency.

## Notes

### Competing Interest Statement

The authors have declared no competing interest.

## References

1. Kosch R, Delarocque J, Claus P, Becker SC, Jung K. Gene expression profiles in neurological tissues during West Nile virus infection: a critical meta-analysis. BMC Genomics. 2018 Jul 13;19(1):530. doi: 10.1186/s12864-018-4914-4. PMID: 30001706; PMCID: PMC6044103.

2. Mei B, Ding X, Xu HZ, Wang MT. Global gene expression changes in human peripheral blood after H7N9 infection. Gene. 2014 Nov 10;551(2):255–60. doi: 10.1016/j.gene.2014.08.062. Epub 2014 Sep 2. PMID: 25192803.

3. de Kruif MD, Setiati TE, Mairuhu AT, Koraka P, Aberson HA, Spek CA, Osterhaus AD, Reitsma PH, Brandjes DP, Soemantri A, van Gorp EC. Differential gene expression changes in children with severe dengue virus infections. PLoS Negl Trop Dis. 2008 Apr 9;2(4):e215. doi: 10.1371/journal.pntd.0000215. PMID: 18398488; PMCID: PMC2274954.

4. Tiwari SK, Dang J, Qin Y, Lichinchi G, Bansal V, Rana TM. Zika virus infection reprograms global transcription of host cells to allow sustained infection. Emerg Microbes Infect. 2017 Apr 26;6(4):e24. doi: 10.1038/emi.2017.9. PMID: 28442752; PMCID: PMC5457678.

5. Cumberworth SL, Clark JJ, Kohl A, Donald CL. Inhibition of type I interferon induction and signalling by mosquito-borne flaviviruses. Cell Microbiol. 2017 May;19(5):e12737. doi: 10.1111/cmi.12737. Epub 2017 Mar 22. PMID: 28273394; PMCID: PMC5413821.

6. Angleró-Rodríguez YI, Pantoja P, Sariol CA. Dengue virus subverts the interferon induction pathway via NS2B/3 protease-IκB kinase epsilon interaction. Clin Vaccine Immunol. 2014 Jan;21(1):29–38. doi: 10.1128/CVI.00500-13. Epub 2013 Oct 30. PMID: 24173023; PMCID: PMC3910921.

7. Zhao LJ, He SF, Liu Y, Zhao P, Bian ZQ, Qi ZT. Inhibition of STAT Pathway Impairs Anti-Hepatitis C Virus Effect of Interferon Alpha. Cell Physiol Biochem. 2016;40(1-2):77–90. doi: 10.1159/000452526. Epub 2016 Nov 18. PMID: 27855377.

8. Matlin, A., Clark, F. & Smith, C. Understanding alternative splicing: towards a cellular code. Nat Rev Mol Cell Biol 6, 386–398 (2005). 10.1038/nrm1645

9. Wright, C.J., Smith, C.W.J. & Jiggins, C.D. Alternative splicing as a source of phenotypic diversity. Nat Rev Genet 23, 697–710 (2022). 10.1038/s41576-022-00514-4

10. Kornblihtt, A., Schor, I., Alló, M. et al. Alternative splicing: a pivotal step between eukaryotic transcription and translation. Nat Rev Mol Cell Biol 14, 153–165 (2013). 10.1038/nrm3525.

11. Baralle, F., Giudice, J. Alternative splicing as a regulator of development and tissue identity. Nat Rev Mol Cell Biol 18, 437–451 (2017). 10.1038/nrm.2017.27

12. Park JW, Fu S, Huang B, Xu RH. Alternative splicing in mesenchymal stem cell differentiation. Stem Cells. 2020 Oct 1;38(10):1229–1240. doi: 10.1002/stem.3248. Epub 2020 Jul 15. PMID: 32627865; PMCID: PMC7586970.

13. Choi S, Cho N, Kim KK. The implications of alternative pre-mRNA splicing in cell signal transduction. Exp Mol Med. 2023 Apr;55(4):755–766. doi: 10.1038/s12276-023-00981-7. Epub 2023 Apr 3. PMID: 37009804; PMCID: PMC10167241.

14. Blake D, Radens CM, Ferretti MB, Gazzara MR, Lynch KW. Alternative splicing of apoptosis genes promotes human T cell survival. Elife. 2022 Oct 20;11:e80953. doi: 10.7554/eLife.80953. PMID: 36264057; PMCID: PMC9625086.

15. Liao KC, Garcia-Blanco MA. Role of Alternative Splicing in Regulating Host Response to Viral Infection. Cells. 2021 Jul 8;10(7):1720. doi: 10.3390/cells10071720. PMID: 34359890; PMCID: PMC8306335.

16. Pozzi B, Bragado L, Mammi P, Torti MF, Gaioli N, Gebhard LG, García Solá ME, Vaz-Drago R, Iglesias NG, García CC, Gamarnik AV, Srebrow A. Dengue virus targets RBM10 deregulating host cell splicing and innate immune response. Nucleic Acids Res. 2020 Jul 9;48(12):6824–6838. doi: 10.1093/nar/gkaa340. PMID: 32432721; PMCID: PMC7337517.

17. Lee Y, Rio DC. Mechanisms and Regulation of Alternative Pre-mRNA Splicing. Annu Rev Biochem. 2015;84:291–323. doi: 10.1146/annurev-biochem-060614-034316. Epub 2015 Mar 12. PMID: 25784052; PMCID: PMC4526142.

18. Ule J, Blencowe BJ. Alternative Splicing Regulatory Networks: Functions, Mechanisms, and Evolution. Mol Cell. 2019 Oct 17;76(2):329–345. doi: 10.1016/j.molcel.2019.09.017. PMID: 31626751.

19. Lin S, Fu XD. SR proteins and related factors in alternative splicing. Adv Exp Med Biol. 2007;623:107–22. doi: 10.1007/978-0-387-77374-2_7. PMID: 18380343.

20. Sahebi M, Hanafi MM, van Wijnen AJ, Azizi P, Abiri R, Ashkani S, Taheri S. Towards understanding pre-mRNA splicing mechanisms and the role of SR proteins. Gene. 2016 Aug 10;587(2):107–19. doi: 10.1016/j.gene.2016.04.057. Epub 2016 May 3. PMID: 27154819.

21. Martinez-Contreras R, Cloutier P, Shkreta L, Fisette JF, Revil T, Chabot B. hnRNP proteins and splicing control. Adv Exp Med Biol. 2007;623:123–47. doi: 10.1007/978-0-387-77374-2_8. PMID: 18380344.

22. Chen, M., Manley, J. Mechanisms of alternative splicing regulation: insights from molecular and genomics approaches. Nat Rev Mol Cell Biol 10, 741–754 (2009). 10.1038/nrm2777

23. Sehrawat S, Garcia-Blanco MA. RNA virus infections and their effect on host alternative splicing. Antiviral Res. 2023 Feb;210:105503. doi: 10.1016/j.antiviral.2022.105503. Epub 2022 Dec 23. PMID: 36572191; PMCID: PMC9852092.

24. Boudreault S, Roy P, Lemay G, Bisaillon M. Viral modulation of cellular RNA alternative splicing: A new key player in virus-host interactions? Wiley Interdiscip Rev RNA. 2019 Sep;10(5):e1543. doi: 10.1002/wrna.1543. Epub 2019 Apr 29. PMID: 31034770; PMCID: PMC6767064.

25. Ashraf U, Benoit-Pilven C, Navratil V, Ligneau C, Fournier G, Munier S, Sismeiro O, Coppée JY, Lacroix V, Naffakh N. Influenza virus infection induces widespread alterations of host cell splicing. NAR Genom Bioinform. 2020 Nov 21;2(4):lqaa095. doi: 10.1093/nargab/lqaa095. PMID: 33575639; PMCID: PMC7680258.

26. Maldarelli F, Xiang C, Chamoun G, Zeichner SL. The expression of the essential nuclear splicing factor SC35 is altered by human immunodeficiency virus infection. Virus Res. 1998 Jan;53(1):39–51. doi: 10.1016/s0168-1702(97)00130-5. PMID: 9617768.

27. Dowling D, Nasr-Esfahani S, Tan CH, O’Brien K, Howard JL, Jans DA, Purcell DF, Stoltzfus CM, Sonza S. HIV-1 infection induces changes in expression of cellular splicing factors that regulate alternative viral splicing and virus production in macrophages. Retrovirology. 2008 Feb 4;5:18. doi: 10.1186/1742-4690-5-18. PMID: 18241354; PMCID: PMC2267807.

28. Lin JC, Tsao MF, Lin YJ. Differential Impacts of Alternative Splicing Networks on Apoptosis. Int J Mol Sci. 2016 Dec 14;17(12):2097. doi: 10.3390/ijms17122097. PMID: 27983653; PMCID: PMC5187897.

29. Paronetto MP, Passacantilli I, Sette C. Alternative splicing and cell survival: from tissue homeostasis to disease. Cell Death Differ. 2016 Dec;23(12):1919–1929. doi: 10.1038/cdd.2016.91. Epub 2016 Sep 30. PMID: 27689872; PMCID: PMC5136496.

30. Schwerk C, Schulze-Osthoff K. Regulation of apoptosis by alternative pre-mRNA splicing. Mol Cell. 2005 Jul 1;19(1):1–13. doi: 10.1016/j.molcel.2005.05.026. PMID: 15989960.

31. Akgul C, Moulding DA, Edwards SW. Alternative splicing of Bcl-2-related genes: functional consequences and potential therapeutic applications. Cell Mol Life Sci. 2004 Sep;61(17):2189–99. doi: 10.1007/s00018-004-4001-7. PMID: 15338051; PMCID: PMC11138917.

32. .Menghani S, Chikhale R, Raval A, Wadibhasme P, Khedekar P. Chandipura Virus: an emerging tropical pathogen. Acta Trop. 2012 Oct;124(1):1–14. doi: 10.1016/j.actatropica.2012.06.001. Epub 2012 Jun 18. PMID: 22721825.

33. Mishra AC. Chandipura encephalitis: a newly recognized disease of public health importance. In: Mishra AC, editor. National Institute of Virology Commemorative Compendium. Golden Jubilee Publications, Pune, India: NIV; 2004. p. 1–20

34. Bhatt PN, Rodrigues FM. Chandipura: a new Arbovirus isolated in India from patients with febrile illness. Indian J Med Res. 1967 Dec;55(12):1295–305. PMID: 4970067.

35. Rao BL, Basu A, Wairagkar NS, Gore MM, Arankalle VA, Thakare JP, et al. A large outbreak of acute encephalitis with high fatality rate in children in Andhra Pradesh, India, in 2003, associated with Chandipura virus. Lancet 2004; 364 : 869–74.

36. Gurav YK, Tandale BV, Jadi RS, Gunjikar RS, Tikute SS, et al. (2010) Chandipura virus encephalitis outbreak among children in Nagpur division, Maharashtra, 2007. Indian J Med Res 132: 395–399.

37. Tandale, BV., Tikute, SS., Arankalle, VA., Sathe, PS. and Joshi, MV. 2008. Chandipura virus: a major cause of acute encephalitis in children in North Telangana, Andhra Pradesh, India. J. Med. Virol., 80: 118–124.

38. Chadha MS, Arankalle VA, Jadi RS, Joshi MV and Thakare JP. 2005. An outbreak of Chandipura virus encephalitis in the eastern districts of Gujarat state, India. Am. J. Trop. Med. Hyg. 73: 566–570.

39. Dibyakanti Mandal. Chandipura virus replication in Vero cells is inhibited by host restriction factors APOBEC3G and IFITM3. The Microbe. June 2025. 10.1016/j.microb.2025.100346.

40. Sreejith Rajasekharan, Jyoti Rana, Sahil Gulati, Sanjeev K. Sharma, Vandana Gupta, Sanjay Gupta, Predicting the host protein interactors of *Chandipura virus* using a structural similarity–based approach, *Pathogens and Disease*, Volume 69, Issue 1, October 2013, Pages 29–35, 10.1111/2049-632X.12064

41. Ghosh S, Dutta K, Basu A. Chandipura virus induces neuronal death through Fas-mediated extrinsic apoptotic pathway. J Virol. 2013 Nov;87(22):12398–406. doi: 10.1128/JVI.01864-13. Epub 2013 Sep 11. PMID: 24027318; PMCID: PMC3807914.

42. Meshorer E, Bryk B, Toiber D, Cohen J, Podoly E, Dori A, Soreq H. SC35 promotes sustainable stress-induced alternative splicing of neuronal acetylcholinesterase mRNA. Mol Psychiatry. 2005 Nov;10(11):985–97. doi: 10.1038/sj.mp.4001735. PMID: 16116489.

43. Livak KJ, Schmittgen TD. Analysis of relative gene expression data using real-time quantitative PCR and the 2(-Delta Delta C(T)) Method. Methods. 2001 Dec;25(4):402–8. doi: 10.1006/meth.2001.1262. PMID: 11846609.

44. Choi N, Jang HN, Oh J, Ha J, Park H, Zheng X, Lee S, Shen H. SRSF6 Regulates the Alternative Splicing of the Apoptotic Fas Gene by Targeting a Novel RNA Sequence. Cancers (Basel). 2022 Apr 14;14(8):1990. doi: 10.3390/cancers14081990. PMID: 35454897; PMCID: PMC9025165.).

45. Juan-Mateu J, Alvelos MI, Turatsinze JV, Villate O, Lizarraga-Mollinedo E, Grieco FA, Marroquí L, Bugliani M, Marchetti P, Eizirik DL. SRp55 Regulates a Splicing Network That Controls Human Pancreatic β-Cell Function and Survival. Diabetes. 2018 Mar;67(3):423–436. doi: 10.2337/db17-0736. Epub 2017 Dec 15. PMID: 29246973; PMCID: PMC5828453.

46. Xueyue Wang, Jiaqi Wang, Yuda Cheng, Jiali Wang, Yanping Tian, Junlei Zhang, Jiaxiang Xiong, Yi Yang, Wei Wu, Ping He, Gaoke Liu, Yixiao Xu, Lianlian Liu, Bangqi Ren, Yan Ruan, Rui Jian, Identification and functional comparison of Bcl2 splicing isoforms in mouse embryonic stem cells. Biochemical and Biophysical Research Communications, 2020 : 524(2), 502–509.

47. Änkö ML, Müller-McNicoll M, Brandl H, Curk T, Gorup C, Henry I, Ule J, Neugebauer KM. The RNA-binding landscapes of two SR proteins reveal unique functions and binding to diverse RNA classes. Genome Biol. 2012;13(3):R17. doi: 10.1186/gb-2012-13-3-r17. PMID: 22436691; PMCID: PMC3439968.

48. Jensen MA, Wilkinson JE, Krainer AR. Splicing factor SRSF6 promotes hyperplasia of sensitized skin. Nat Struct Mol Biol. 2014 Feb;21(2):189–97. doi: 10.1038/nsmb.2756. Epub 2014 Jan 19. PMID: 24440982; PMCID: PMC4118672.

49. Suri P, Badalov A, Ruggiu M. Alternative Splicing as a Modulator of the Interferon-Gamma Pathway. Cancers (Basel). 2025 Feb 10;17(4):594. doi: 10.3390/cancers17040594. PMID: 40002189; PMCID: PMC11853465.

