## Supplemental figure 1 for "Differential modulation of host cell Serine and Arginine-Rich (SR) protein and apoptotic factor BCL-2 gene expressions during Chandipura and Dengue virus infections"

Supplementary Figure 1 : SRp55 binding motifs in apoptotic factor BCL2 gene. Mtof sequences (TCCGGA) are highlighted in blue color. Exons are in uppercase and introns are in lowercase letter.

CTCGATCCGGGCGATGGAGGAGGAAGCAAGCGAGGGGGCTGGTTCCTGAG

CTTCGCAATTCCTGTGTCGCCTTCTGGGCTCCCAGCCTGCCGGGTCGCAT

GATCCCTCCGGCCGGAGCTGGTTTTTTTGCCAGCCACCGCGAGGCCGGCT

GAGTTACCGGCATCCCCGCAGCCACCTCCTCTCCCGACCTGTGATACAAA

AGATCTTCCGGGGGCTGCACCTGCCTGCCTTTGCCTAAGGCGGATTTGAA

Tgtaggtggtgcgggggagcgggagtggggcgggggggactgcccaggga

gtgactttccgaggaaggcatttcggagaagacgggggtagaaaaggctg

gtgggagattcagagtccactggtgctttcgatttgacttaagtgaagta

tcttggaacctagacccagaccttcgtaagacccacaaagaaaccagttc

tggtacctggagggggaatggaatttttagggtaaatggcatgcatatta

attatttttttttcctgaag CTCTTTCTCTCCCTTCAGAATCTTATCTTG

GCTTTGGATCTTAGAAGAGAATCACTAACCAGAGACGAGACTCAGTGAGT

GAGCAGGTGTTTTGGACAATGGACTGGTTGAGCCCATCCCTATTATAAAA

ATGTCTCAGAGCAACCGGGAGCTGGTGGTTGACTTTCTCTCCTACAAGCT

TTCCCAGAAAGGATACAGCTGGAGTCAGTTTAGTGATGTGGAAGAGAACA

GGACTGAGGCCCCAGAAGGGACTGAATCGGAGATGGAGACCCCCAGTGCC

ATCAATGGCAACCCATCCTGGCACCTGGCAGACAGCCCCGCGGTGAATGG

AGCCACTGGCCACAGCAGCAGTTTGGATGCCCGGGAGGTGATCCCCATGG

CAGCAGTAAAGCAAGCGCTGAGGGAGGCAGGCGACGAGTTTGAACTGCGG

TACCGGCGGGCATTCAGTGACCTGACATCCCAGCTCCACATCACCCCAGG

GACAGCATATCAGAGCTTTGAACAGGTAGTGAATGAACTCTTCCGGGATG

GGGTAAACTGGGGTCGCATTGTGGCCTTTTTCTCCTTCGGCGGGGCACTG

TGCGTGGAAAGCGTAGACAAGGAGATGCAGGTATTGGTGAGTCGGATCGC

AGCTTGGATGGCCACTTACCTGAATGACCACCTAGAGCCTTGGATCCAGG

AGAACGGCGGCTGG gtaagaaccaagccccttgtgtgtcccttttctttg

gcctctggtcagagatccccaacagcccttcttctgtatctctttctgtt

gtgttgggtgattgttggagacgttgatagttgaggaaacctgactggcc

tcatttcaccacaagaggttaactccttggaaatacaaatgaccctgtgc

ttcagagatcaggcatctttcaccctggtcacccccatgactccactatt

ccatacttgtgttccagtttctcagcaaagaaaactgtagcctatgtgtt

cattaagtcccaaacctgtgagggtaggaagttagttcggtgtttccctt

accttatacaatggttcacacacaaaggggtgatgagtataatgcctgga

gaagatgatgagcatcaatgacccacagacctctccatctgttcccccca

aggggtctcagtcctatcctgggcttctctcccccaaatcaagttccctt

tatttcaaagtttgcgtgtatggcactctcctctcagactagggcatggc

cataggcaggagttcccactgggttgaatggctgagaattgtcagatttt

ggggcagtgtaggctgtgcagattgtgtttgcactgcgccccctaagttc

atttcccccaaagtgtgcgactttctgcatggttgccttgtttgtggagc

acttgctgtgtcacacattaagccaggggactttgcacctcattgcctgg

ggtcggttggggagctccagctgctctgcctgttactttccggccagagc

ccaggggcctcttcaagcactgtttgctggcaaagcaagaacctagtctt

tggggctctgtgggacattgagatttagtaacagacttcacatgcaggta

taagttccagttctgtcatgtaactagctgtgtgaccttggacaggctgg

gttactttcttgaggcttagtttctttcactaaaaaatgtggcaagctct

ccttgccttacagtgtattgtaaatcttggcgataatgtgagcacagtga

ctgacaccagcaaattccctttccagggagggagagttctcctgcccatc

actgagccaggatgagatggtggtgaaagcttctttaccagtggtcactg

taggcaagttttgctcttcagtgaattcagtcctgttttcttgcttatct

ccctacctagcggaggtgcaggttttatgagagatgttctatcccctcag

agtatgcagaaaagcatcaagctgtggaaaataagtcatccattgaagga

gctgaggtggaaggggacatgcaccatggactcaggaatgtcaagaatgc

aactgtctataggctgggcctagggtgtggcaggcacttcactttaccac

tgagcaagagacctcctgagtttgccctggaaccccaccccctctctcat

tttgcatttctttgcaatttcaactcgtctgtgccagctcttgtaaaagg

caagggggtcaagtcaatagaggtcagataactggggtggggtcagggga

ggtaaaggtggtgggttctcttgagctagctgactagacttgaaagccat

tctgtaaatgaactctctagggggataaatgattaaatagttcctgttgg

tttgtgcagtgacctctcctgttccctcctgctttgattctcttttgggg

caaatatttgttcagctgattacaggagggacaggattccagtactgcag

acacaccctctgaggcatctccaagtagggattgaggctgcttgcccact

ttgcctgactattgtctttggccagtggtcctaataccaggtgggttgtt

gccagctataaacaaaaggcagtctgggagaggtgcctggtctgctagcc

agcctctgcagaaggctacccctaccagagaattgccagctgcccagtgg

tgcatatgaaagaggtcttagacacagcaactctgggtaagtttattaga

tctccatggggttgaaattccatcttccatctctttgcttaagaatcatc

agagcctgttggacaacataccacatgtcacttcctcttggtgaacagag

gggatgtgtgtggaagagccagttatatcagaaccatttctgagcacaca

ttctctccctaaggccagggagtaaagaacatttgactcaggatcctttg

agcctgggtagaaggtacccgcccccttcctctcaggtattatcctgttc

caagctgcaccgactggctactgtggtggcatcgggtaattatcctggag

actgcggctgggagatccttaagggtaggagttatgggctattctactca

gggctatgttctaaatttagcatccctacaggaaaggggaaggaaattag

tctttattgagtacctgttaagtactaggcactgtgttaggtcctctact

tatgcaaccagtgtttcttgagcatctactgtgaagctagcagtgtacta

ggtgctggccataagccacgcatggtaccttccttcatggaaacggcatg

ctgtttcatatattctacagtaaccttccaaggaaggtgagtttattccc

agtttacacatgaaggtctttgtgcctaagtgactgaccccaggtgtccc

tcagccagttaagtgtctgaatgagaatttacaccaaggttggcataatt

ctaaagccaacacttttcatcctatcaagttgtccagggagtaataaagt

gacctagacatgctggcccagaaagccagagatgctgttgggccaagcaa

accatgcctgtgaaggtgctcactctggactgataactggctttgttcag

gtagctatgggaaggatagttggcagtagcaggttcccagaggcgtgttg

gtagcctcagattctggcactggtaaaactcagacatggcaggtttcaag

tgggcatcagggttttgcccagagtgagctgcttattaattttttctttt

tttttttttttttgagacggagtctcgctcttgttgcctaggctggagtg

cagtggcgcgatctcagctcactgcagcctctgcctcccgggttcaagtg

attctcctgcctcagcctcccaagtagctgggattacaggcatgcgccac

cacacctggctaattttgtatttttagtagagatggggtttctccatgtt

ggtcaagctggtttcaaactcccaacctcaggtgatccgcctgcctcagc

ctcccaaaatgctgggattacaggcatgaaccaccgtgcctggccccggc

caattttttcattgttaaaaccctctcaggccctgccaatagatgacagt

ggtgtgcgtgtggggagaaggtggcctgccccaggtatctctacttagtg

gtggagtgatgcttcaaggactgaaagtgtagacccttagccagattact

gcctgcctggagtcacctgatgaaatgctgaacctgttcggctgtcttgt

ccatctgtctctcccgcctgtaaaatggaataatgatagccagctgccat

agcttgactcgagtaaaggatctagcacaaaagcccttctcctgccataa

ttgtagggtggaaaccctttaatgaatgggcagcttccctcagagaactg

actcttccccctagagtttctccttgagaaacaaagtccctgtgatactt

tcctggaatgttgtatacatgaccttccccgaagggacacaagtgtttct

ggtgctttccaatgggaatgtgggaagggacccaggtgggccttgccact

ttgggattgctgtccctgaagaaatcccttagcctgatagaaacgtaatt

gttgggagcaatgaactgtgttgggggagaaaacataacttggcctttct

taagctgtatggctcagtggtctgagtttctgtagatctcttattgccta

atgtggcttcttcagatggatgtatgcttagttgaagcacctcagctccc

tttgcagatagtggcagttctgtggttttcttagggaaaccccattagaa

accccaaatgtggaaatttttagtaatctcattcgggtcccaaaattaag

tagaagtgtctcaggggctcccccttgaaacacctacttcttctgagtgt

acatttgaggttgctcttaaaattctgttaccatcacagggttttccaga

cttcctaggaataatttccccagtagccccacagagccctttagcttctg

tcgaaaattacccgggtattttttcagggggctccaatggaacctggagg

gttttcttgacttttgtatgggtgtccccagatttgtttgggctgcatga

aacagcagttgttatctctgtcccttcccccagcagatatctgtcctgag

gccagtgggcttgagcgtcctcctctgtgtccagtctccctggttttgcc

agtgccctcagccttccatcatggggctttcttgaagcaactgtggtttt

gcctgagtttaatgtggatagacataggggtaattcccagtgtactgctt

ttctacggaagtataatctcctactgccttaattaagcagaggccctgtc

cccacccaacctccaacctaattccacagagtcctccctaaggactccct

cttttttctttgaaggagtcactctactcctggatcctcttctcttctgc

ccctctatccgacagtctgtcttccttgttcccagttggcagacctgagt

caggcaatcccactgggtctttttgtttagcctggggaagagaagaccca

ggagaatatcaccagcttcagatatctgaaggaagagggactgagtcttt

ttttattggtgtggtacctggcaaaacttggataaatgaaaagaagttaa

cagttggtagagcttaaattatgtgcagagaagacatttctaacagattt

actctgtcctagatttaggagctcctcagggtggaggccacatcttgtcc

atgcttgtttcctcagcactgagcagaggccttgggcatagcagatacct

aattgaagagaaacccactaaatttctgaagaaacagacccttgtcaagc

cagaataacaataatgggagtggctatttacctaataattcattcgttga

gtcaaacaatgtttattgagctcctactgtgtaccatactgggagtatgg

cagtaaacaaaagtaagcaaaactaagtttctgtcctcatggagctcaca

ttccaatgaggagtgacaggcaataaatatataattacatgtttatatca

gatgattgtacatgctatgaagaaaataaagcaggttggggcatagggca

tgtctagcaagggagggtggggctgcagttttaaaagagtgatcaaagaa

ggtgtaagggagaaaaaaatggctttcctctacccttctaggtcctttgg

ctgggctgagaattaaattgacatagacagattaactggaggaaaaataa

ttaattacatagttgtacacaggaatcctacaaaatatgaggctcaaaga

agggtcagataattgaagcttatatagcatcctgagctacagaaaagaat

atggggcttgggacttctgttaggggcttgagtggcagcaacccaagtta

tggcagggtgagggaaagaattggtggtggcgtgtgtctgtggtcccaga

tactgttgagggggtgtactgaggtgggaggatcacttgagcctgggagg

tggaggttgcagtgagccgagatcaagccactgcgctccaacctgggtga

cagagcaagaccctgtctcaaaaaataaaaataaaaagtctctctggtaa

taaaatttgtctggagcagccctcttcctaatacagatacctttactagt

gtagatttcctttttagatgtgaatttcttttacaaaaggacagcttttc

agagctactcttatgtctgcagtttctcagaataaccagttcaaaatatg

ccaaagaagtatatttttggggtggcatattgttttcctacagtcatatt

ttgggatggtatgtcctgagtcccaacaaaagtattacctaaatttattt

gagccaagcgctaaatgaggtaaggaggtggatgatgtaggtaccctaca

gcaaagggtgtttaaagcagagggaatggactcaggagggagcatgcttg

acattttcaaggatgagcaagaaggtcaatgtggctggagaagaaacagc

aaaggggaaaagggtagaaggtggggagggaatggttgaggaggaggatg

gagaagtaggtgggggccagatcgcatacagcattacagaccatgatgag

gatttggctttggtggaatggtctgtattccttttttttttttttttgag

acagagtctttctctgtctcccagactggagtgcagtggcacgatctctg

ctcactgcaagctgcacctcctgggttcacgccattctcctgcctcagcc

tcccaagtagctgggactgcaggtgcccgccaccacacccggctaatttt

tggtatttttagtagacggggtttcaccatgttagtcaggatggtctcga

tctcctgacctcgtgatccgcccgcctcagcctcctaaagtgctggaatt

acaggcgtgagccaccgtgccaggcctggtctgtattcttaagggtggca

acagggagcccagtgaagaggctggtgcaattgccaggtgaaaaagaata

gtggcttggacctagaggtggaggtgtagtgagacggggactggagaata

agaccaggcttgcacatggtatattgatgggtccagccatcctggaaacc

atatttttcaactacttggaggtcctagaatcaaagttaagtaagataga

tgttctttgggaatagttctctccaaatgtgatattctgatgaatggtcc

tcttccactccaacagtatttctgtctgagtaaatgaaatttgtctgagt

tcagttcccaactcatccgtgtctacctatgcaagctccttttacctctc

tgaatcttgggacaaataataatacctgcctcacagggggcgatggtaaa

gacacagtgaggtatacctgagatgcataaagactggagccatgttggta

tcattattcttattatctttccacagaacgcatgtcagagacccattcac

ggaatcttgagcatccactctcctttttctaaataccatagtgacccatc

agcacatctcagagcaggcccttgcagcatttctgttgtagtaggagctc

cacagtgtttgtgtgcaatggtggtgtaagcactggtcttggagttggaa

tcccagttctactctcagtctcaccttggccaagtcacttcacccccagt

ccccagtttccctttatgcaagatggagggtttgaagtaggcaataaagg

aatgctggttctctctatccttaggccttgtggcggattccaccattaca

gcatatttcacattagttatattttaaaatggctctctcctgttgacaga

tatgtcagcaccttaactaaacccagtgtcaaggcaggctagtttatcac

tgagcccacaatagtgctaccctatatttatgtaacaccttacagcttcc

aaagcactttcagatccaccttgcctttttgatccttataaccacccaga

gaggtgagcaggaaggatgacatctccgtttcaaaggagagaaatcaggt

gagaaagatgaattgacttgcccaaggttacatcaggaggaagaggcagt

ttttggactttcaaattggaacttgactactagcttgctgtgtgaccttg

ggcaagtcttttaacttgtaaatggctatagtaatgccacctttttctga

agattgggtgaattaattcattcttttaaggctttcatatcatcctaggg

ctcttcactcacatcatctggcttctctttttttacatattgatttttga

aattgtggttttgtataaagaaactgcctttgggaccgtctgctatttgg

tttcatccctgatgagactctcgggccaagcttgagccttacagtgaact

ctgagccacacagctttttgggcagcctccagtgctcccctcggaactgg

aggtttttctccagctcttcaggaaatgccattgcgatgacttccatgga

ctgcttcctgtcccagccctgccggcaagagtgtggaaactggaaagcaa

tgtccctagaggtccgccccccaaaaactactttttttggacgtgtccct

gcctttcctcttgtccattactttctttgctccctaccctttcctccttc

acctggcttctgcctccagctgtgatcatcccttcaagatttagaaaatg

cctttctacttggttctgccaggccattaccccctaccctaggctaggct

gtgagggaggtcagttagaattcctgggttctctccttggctccatcacg

cattagcagtttcagtcctggacttcccctggagatgcattaaacctgtt

gcttgcggtgccctgtgtcatccattaaccctgagttcccctaggagatt

ccaccccacccctgcataggatgggattcttaagagataacttgaggttt

ggcacatataaatcctgcctcttggaaattgatgataaagggtggagtga

aacagagcctgccttcccctcccccagtcacctttgaggaggaagctaga

aggttgaaaaccttgttggacaatcatctgaggaatgggagctggcctgg

gaggggattggaacgcactcaactatgcctcagatcctttgttcctctct

tgaaagggaaacattcccctccctcctccctggccacacacacatacaca

cgtgcctttgttcccccgagtcaggcagggataggaacatagtgtccctt

ccaaggagcctgactcgcttccctagagactggcctcctggtcagtggga

ctgggatatttgctcaggagcccccagcctcgtgtcctttccatagagtc

ttcgaggctaaaaacttgcctacaaacagctgctgagacttctctgtaat

tatttcatgggacacccatgtcccaggataaacagtccctccctcttact

ccagttggtgattaatattattgttaacattactattaatagtatctaac

acttattgagtgcttactctatgccagccattgttcaaagtgctttacat

gtattaatagatttcattctcacagcagcactctgaagtaggttctttta

tccctgtttcacacatgaggaaactgaggtacagagaggttaagcaattt

atgcaaagtagcacagttagtaagaggcaaagcctgggagggtagcttcc

aacctcccccttttaaacgctccactgtgttccctcgtccagtgcgctat

tgtctaggtgactgctggcattaatccagtgctcactatgtgacagcctt

tagtgacgctgtttttgttactatcccaacaaccttatcgttataggcac

tgagatttgggagggcttatagttagccagaggcagagtcagcatttgaa

accaggtctctcttactctgaagcctgtcctttttatctccagtccttaa

aaatcttgcaaagtaggtgttgttaccttcattttatggttgagaaaatt

gaggctcagagaaaagtgatatacccagagcttcacaggctagtaccaag

aagagctgggattcacactctactgtcttagtagtctgtattgtcatgat

tattctttttatgtgcatgataatactttcaatttaccaaactcctttta

taagttatttcccagtgaagtaggagaggtgggtatgttcagccctgttt

ccagaggggaaactaagataaggggtcttgtcccacagtaaatggcagca

tcagacttagaattcatgttctggcttttaacacagtattatcaatactg

tctgcctctcctaaagctggatgtgtggcttggagctgtccttggagttt

catttcttttgttaaatcagcaaatattgacaaatgctttgtaatgtgct

aaatgctagtgatactgtggatattgtggtaactgagactgagtcccatc

cctcaatggtctgtttccatactccacaccttgggtcttgctggtgtccc

tctgccatctgaatcagtctacatgtgccatccatccttctcctcagctg

ccagacacaccaggcaagattcattatagttgatatgtattaatcacctt

caacacatacacagacaaaactctggcctcacatttgagacagcttgact

ccttgctggtggcataacacaaagcaaatcagcttcaaattctcatccat

aagcctgagagaataatgccaagccaccagagtaagtgaggcatgaatga

gaaagtgaaatgtaagtgtgttttttggtaataataacagatactatggg

ttgagtagttacttgctctgtgccaggtgctgtgtttatcccctcaatat

aatcatcttacttggttctcacaaatgctctcctgtgaagatggagaatc

tagggttcagagaggctccctaatgtgtccacagtaacagatcttctaag

aggcagagctgggtcttatactcagatgtgtctgatgcaaaagcctataa

gcgtcattgcttcttacaagcccagactggcttattttaccctgactacc

tggctcagggcccgcggttgggtgaggactcagccagggcctgctcaatg

aacatgaagggttattattagatacaggttaggctacaaaaaaagtttct

gcccctaaggaagcaaactcatttgaagcagtaggaatagacctcaaaca

cttggtgaatggcttgaggcaaactgtgccaaacctgtagagtaaacacc

aaaagtggaagaagttgagcccacctgactgcattgctgggccctgggcc

tctgccaaatgggccagcttccaccatcagctacccagttgattactggg

cccagaggcctccaccctcagcctgtatcatcaacctgcaccctttccct

agccaactgcattgccaatccagactgtcttctgcatcaacagtggaaag

ggttaggaaggagagaggtcactgacagcttaaataatcatagctttctc

agcaccgtagctttgtacaatttccactgaggcctcagtttcccggtctg

taaatggagataatactaatctcttaatgaagatgatagtaatctcttcc

ttataggcttgctgtttttcctctgcctgtatcactcatcctatggatct

gatcagggctctctctcttcttatcattcaggtatcaactcagatatcac

ctccctagaggtcgccatgtttacatgtttaccatctgtctcctgtagta

aaacataagctctcggagtacacgggctttatcctgtctttcatccccag

catctagcagagtgcttggtacataggaggcacttagcaaatatttgttg

aataagtgactgtgtagaccacctcacacggtgtctgatatactcagtca

atgttaggcattgttatgtcttaggaagatgtggaccatgatgggtttaa

ataggtgaaaagaatgtaacaagagcaggctgtggctgggcgtggtggct

cacacctgtaacctcagcactttgggagactgaggcaggtggatcacttg

agatcaggagttcgaggctggcctggccaatatgacaaaaccttgtctct

tcttttttttttttttttttttttttttgagatggagtcttgctcttgct

cagttgcccaggctggagtgcagtggcacgatctcggctcactgcaagct

ccgcctcccaggttgacgccattctcctgcctcagcctcccgggtagctg

ggactacaggtgcccgccaccaggcccggctaattttttttgtattttta

gtagagacggggtttcaccgtgttagccaggatggtctcgatctcctgac

ctcgtgatccacccgcctcagcctcccaaagcactgggattacaggcatg

agccaccgcacccggctcgtctcttctaaaaatacaaaaattagcagggt

gtggtggcgggcgcctgtaatcccagctacttgggaggctgagtcaggag

aatcacttgaggccgggaaatggaggttgcagtgagctgaaatcatgcca

ttgcactccagcttgggcgacagagggagactctgccacacacacacaca

ccaaaaagatcaggctagttacacacgcgtgcacgcgcgcacacacacac

acgagcaggccagacgggtaagaaccacccagacagaactggagggtgga

gtatgtatgaagcatttgagagatcatgaggaggcagactggctagagcc

aaagatggtctgtgtcagcaataaagttggtcagggaagaaaccaggtgg

ctgatgacagaggcctctggacagagggatggttgatggtagaggtgggc

ctaatggggagaggcccagagcacctgtgggatgcagttaggtctcagta

acccctaagaaaaagaatctcttgccagccagggctcagcaggcagagct

gcagggagagaaaagccagaggcacctgcaacctgcagatcaggtaagct

gccctggttgttattttgggactcagggccaggtgcagattcaaaggatt

gattgaagccttttggcaactctggcctcagcagggtaaacatcacctgt

ctcctgccacctgcagtgacaatgccaggaagggtaccctaggtaggccc

aaaggggcctaactttcttgctgccctggtggagcaagaaatatcttcta

ggagaggccaatccttccctgccccacagctccttctctgcaaagctcag

gtggcaatcaggtacctcctgcccaagaggcccccatggttctggcctaa

ggaaggcagggcggggcattgggagccgttgacagctgggctcagctggg

gggaggggtcagtttgggagcaggtgcagatttcagggagggtggggcct

aaagggaagtagggatcttggtaggctgcaaaattttcctccccatcccc

cagccacagctggcttctccaggaacctgatgacctagttacaatgctta

gaaattgtcagcttagcttgcagttcagggctttgccagtctctcctctc

tgaggcttcttgggtccaaaaggtaccttggcatttccccaagggggttc

catgtgaatgacatacactgtatcacatgtacaggcttcttgcccagggg

aggaaagagccagtaaggctgctgcttcttcccttgtgtgtgccccccaa

ccccagggagaaaatactcccatgcccttgcccccttctgtttctctcta

gagggttggtgagtgacaaatgggaagtcacaaacttcttcccttgaggg

tcaacaaagagggccagacccttgttgccctaaggatagattgagcacac

ctaaccattctttccttacctccccatctcagtgtgatgttaaaacctga

tggaaagaggttaggaagactcttctgggggctctggaggccccccactc

tctcaggcagcagctgctcctggtattcctggtgtctgccttgcctcctt

gggagaaattccatcgttgttaagagtcaggtacgaaatttgacaaaccg

ttgttgagttggtgagtctcacttgcttaatagttatccttggcaaatac

ttaatctttcccagccttagtttcctcatgtatggaaaagaggtaacaat

atacagtgcttttcttttttccttttttttttttttttttgagacagagt

ttcgctcttgttgcccaggctggagtgcaatggtgccacctcggctcacc

acaaccttcgcctcccaggttcaagtgattctcctgcctcagcctcccga

gtagctgggattacaggcatgcgccaccacgcccagctaattttgttttt

ttagtagagacggggtttctccatgttgatcaggctggtctcgaactcct

gacctcaggtgatctgtccgcctcagcctcccaaactgctaggattacag

gtgtgagccaccatgcccggcctacagcgcttttcttgaaacactcttga

gagcatgaaaatagttgaaaagatacatggagcacttaccaaggagccca

gcatatagtaattacttagtataacaagctactattattactgttattgt

cacattgcaatcactggtaaacaaattgttgtcattgctattggatctac

tctccaggagtgaactttcaggctgacagaatatgtggatatctttggac

caagagtttagagattttaaggtacctgaagcttggggccttggctcaca

accttgtcccctacctcctctacaaggcccaagtcataccttgtacacag

tagaggtttggccagtgttgttgactaggcgtttggcaatggggcatatt

ttgttttatcttcccagggtcctccttgtttgttgactgagtggaagact

taatgagttaagtatttaaagtgggtaggagcatgcctggcacacagtag

gtgcttgacatgggtttgaatgattaaacatggcattgtgggaactctgg

agtttttccaggatggccccactcactccttgtttgctctgtggcctcaa

gcctcttgcccaagctctgcctgttgcctcatatgttgaagaaagataac

acccgccttgcctccctcatgggctagcagggaggcccaaagagatgata

gatgaaaaagcactttgagaactataaatatttggtcttgttgctactcc

ttgactaaagtcacagccatagctgtgtcattcattcattcattcagtca

acaagcattttttgagtagaagcaatatcttatcacttaactggcttcat

gaatttggacaagttacttccctctgtaaacctgaagatctataaagtgg

aagtaatactcatttcagagggttgttgggagaaagactaaacgaggtgc

tctgagcttactgaggataaaaactatgtcttattcactcttaacagcta

ctgctgggcctagtacttggtggctgagtgtttagtgactggattaacaa

acattcctaataattaatggtacatgtaaagtgcacttacacattgccta

gtgcagagtaaaagctcagcaaatgctgagtctttcccttttctctatga

ggcttgaggtgccttgatttagccatgggttgccttctgcagtcaaagta

ccaggcaaggccagactctgaatatgactcagatgggaagtagaagtgag

tattaatgcctggctccaggttgtccttacctgcctacgctggacttcag

tgcgggagttctgaactatcgaggatcagtgctgcttaaacataaaatcc

agtgtactatggttctcaccccatgcttatctctgatccagaacaactgc

ctcttgttctacccactctccatctgctactcagaagaatttccccttgc

tcacagcacccatgttcagttggagccctaggggttaccatgtcagggaa

gtcccagggagtcactcagaaatagcctttggggcaaggtccagagaagg

agcccaattctaaggctcagtgtctcctgggactgggtcaagcagcaggt

gcatacatgagacccagggctggtggctgagtgatgaaactgagagatgt

gtagggccagacaagaagtcaacctagaggaaactgcctgcttgttcatg

caggtgtttggacacctgggggaactttctgggtctgatttagtggtgct

agtgtaggttccatattagtatcccaagccccagtgagggaaacctttgg

gactcccaagcaaagcctgtagttggcacccagcacctttaaacccttct

cagatccctttagggtgaaagagaggccccctcctctgggttctacagca

tctacagctggaaactccagggacaatgacataataaccatagctccctt

tataaagtctctgctgtgtgttaggcactgtgctggggctctttatgaaa

aggggatcattttgttaattggaagtatgggctctggaatcagacagagt

tgagtttgaatctggtctctattacatatcagctgttagattttgggtgc

attcctttccatttctgagcctcagtccccctatctgaaaaatggggata

ataataatacttcatagggtttttgtaaggatttaagagccttttgtaag

aatgtatgtaagatcatatggtgcaggtattaaggtaattttagtaatat

ttaacaatgagcacttgcacttaagtgttttatatgagatagttcattga

attctcaaaatagtaccactcaggtaggttctcttattattcctatagag

aagttaagaaagttgttcaaagatgcacagcgagtaagtggtaaaaccat

gatttgaacccaggtagtctggctccaaagccaacttgtcctacctggat

aaggaaggccctagtaaatgttgctgttataactattgtatccagcattg

tagttgtctgtagtttttggtctatccactgaattctgagctcagtgaag

actagaaacccatttaatcaccttgcacccccagagaccaacataatact

cataaaactcattaatatttgcttaatggctgagaatattctctttcttt

agtaatcaatcagtaactctttatcgaccagttcctgtaggctagctttg

aactaaatcctaggaggaatgcaaagtacagctatgcctgtgtaaacgga

agattgggtgtgtgcgttgggggggtgcattccagagagcactgtgtcat

tggcagaaacttccacacaggaggcagaaacaatacaggtggtttgagga

tgttttccaagatgggattcccacccctccctgagggagtcacttcctcc

atctcctgcagaacaaggtgacacagattccctactccggccaagggcaa

ggaaccactctaagctcttcagtcactctaatccagaaactgtgtcttta

tggctcaggcaaccagttcatggcagcctagaatgacagggaaaaagctg

ggaagggaccttagaaaatcactcttgcccaatactccagccaaagtggt

cttttaaaaaccaagttcaggctgggcgcggtggctcatgcctgtaatct

cagcactttgggaggccgaggtgggtggatcatcaggtcgggagttcaag

accagcctggctaaggtggtggaaaccccgtctctgctaaaaatacaaaa

attagccgggtgtggtgcacgcctgtaatcccagctactcaggaggctga

ggcagaagaatcgcttgaacctgggaggcataggttgcagtgagctgaga

tcgcgccactgcactccagcctaggtgacagggagagactccgtctcaaa

aaaaaaaaaaaaaaggtttagggccgggtgcagtggctcatgcctgtaat

cccagcactttgggaggccaagccgagtggatcaactaaggtcaggagtt

cgataccagcctgaccaatatggcgaaaccccgtctctactaaaattaca

aaaattagccaggtgtggtggcgtgggcctatagtcccggctacttggga

ggctgaggcaggagaatcgcttgaacctgggaggcagaggttgcaatgag

ccaagatcgcaccactgcactccagcctgggtgacagagcaatactcgtc

tcaaaaaaaaaaaaaaaacaagttcagggctgggctcagtggctcatgcc

tataatcccagcactttgggaggccaagacaggcgaatcacctgagatca

ggagttccagaccagcctggccaacacagtgaaaccccatctctactgaa

aataaaaaaattagccgggtgtggtggtgcatgactgtattctcagctac

tcgggaggctgaggcaggagaatcacttgaacccaggaggcggaggttgc

agtgagccgagattgcaccactgcacgccagcctgggcaacagagtgaga

ctcttgtctcaaaaaaaaaaaaaaacaaccaccaagtttaggccaggcat

ggtggctcacacctgtaatcccagcattttgggaggccaaagcaggtgga

tcacctgaggtcaggagttcgagaccagcctggccaacatgatgcaaccc

cgtctctactaaaaatacaaaaaattagccaggagcagtggctcacatct

gtaatcccagcacattgggacaccaaggcgggtgatcacctgaggtaatg

agttcgagaccagcctgaccaacgtggtaaaaccccatctctactaaaaa

tacaaaattagccgggcatggtggctcatgcctgtgatgccagctacctg

ggaggctgaggcaggagaatcacttgaacccaggaggtggaggttgcagt

gagccaagattgcaccattgcactccagcctgggcaacaagagcaaaact

ctgtctcaaaaaatttaataaaataaaaaccaagttcagatcttattgca

tttttgtttacaaccctccaagtggtcttccactgtagttagaataaaac

ccagacttttttttttttttttgagatggagtcttgccctgtcgcccagg

ctggagtgcagtaagtgtcatttcggctcactgcaacctccgcctcccgg

gttcaagcggttctcctgcctcaaactcctgagtagctgggattaaaggt

gcccactggccgggcacgatggctcacccctgtaatcccagcactttggg

aggctgaggcgggtggatcacctgaggttgggggttcgagaccagcctga

ccaacatggagaaacccgtctctactaaaaatacaaaattagccaggctt

ggtggcacatgcctgtaatctcagctactcaggaggctgaggcaggagaa

tcacttgaacctgggaggcggaggttgtgttgagccaagatcgcaccatt

gcactccaccctgggcaacaagagcaaaactccatctcaaaaaataaaaa

taaaaaaataaaggcacctgccatcacgcccagctaatttttgtattttt

agtagagatggggtttcaccatgatgtccaggctggtctcgaactcctga

cctcgtgatctgcccacctcagcctcccaaagtgctgggattacaggtgt

gatccactgcacccagcaaaacccagacttcttaccaacctctgaggcct

gtgtgatctggtcctggctgcccctccatcattctgtctctcccttactc

cactcaagccgcatgggttccttgctgcttctggatcctggcataacttc

ttcctgcccaagggcctttgtagttgctcttttctctgcctgaaatgttc

tgcccctagtatttcctttctgaggcctacatttggttcagcttgctggt

cacctcctctgagagactttccttgaccttcagtttaaagtagccatgct

atttctaaatccctctccagtcttcttgtagttttgtcatagcaacagcc

tctacctggaattttctttttgtttgtgtacttgtttttgtctgtctctt

ctattagaatataagccccatgattatagggcgtttggtttgttcatcat

tgtatctccagtgtctgcagctgttctttgcaggtagtaaacattcagtg

aatactttttgaataaattattaaacaaatgaacaaaacctaagacatga

aaaaactgaggcccagagaagagaagctaatgagaattaatagtgttctt

tcatcctacttttgaatgctcattaagtgccaagcactgtattcagtgat

ttacatgcattatttttgccacttattcttaattgttataatgagttgtt

tggaacatcattttttatatataccactgtaccacttaaaagaaaactgg

accaatcttattttttattggaaccttgctctgttttacaaagagttgga

agcaacttaggaaaattcatacaaccaaaaggataatgtagctaagggag

tcgggtgaaaggaaaatatgggtaagaagacgagccaaaagtaaaattag

ttcacacaaatgtatgttacaaaagtcctttgaactgagcagtttttatg

ccattctttataactaaatattgcccagattcatttaaaatgatggtttt

ttaaaaatctaactacaggccgggcgtggtggctcacgcctgtaatccca

gcactttgggagggcaaggtgggcagatcatgaggtcaggagatcaagac

cgtcttggccaacatggtgaaaccctgtctctgctaaaatacaaaaaatt

agccaggtgtggtggcgcgtgcctgtaatcccagctactcaggaggctga

ggcaggagaatcgcttgaacccaggaggcggaggttgcagtgagctgaga

tcacaccactgcactccggcctggcgacagagcaagactccgtctccaaa

aaaaaaaaatctaattacatagatgaccttgggattgtcccagactgcac

ctaatagtctaggatttgctggctcccttcctgcctcatcagctgggctc

ctggattgaggcaacacgatctcctctttttgccaaaggggctggtccag

cagggaggagcctagacccctctccagtcctctctggacctagctaactt

tgtttcaaggagtcttttctgtagaaggtttattaattgagatattgcca

ttggaaacccagttctgctctcaagatacttccagttgaattagagagaa

aagacataatatacatgaaaggagatcttgtaaaagccaacaataaatgc

taaatgccaaatgagggaaggacttcacagaagaggtgaggtttacaccg

agccttaccacaaagcggagaaaagaggggctgtagctcccagtaggaga

gtcaatgtgggttaagatttgcaggctgagaccaactctcaagaagagac

ccgtgggcagtggcagtcaaattttaggcatatgtggaggggtctctttc

attaggtaataaagccaaaaaggtcagctagaaccaggtttgtaggatct

tcaagtgccagggggaggattccaaggccctccttggcagccctggtatc

ccaatagataggtcgtgcccagggactaatgcccatgctggccactggtt

gggctgctgattcccaggtggcaatggcagcttctgaaatgggagccaga

gccttcaactaatagtgaggggaggagttaagtgtggggccattgtcagt

ttagcttttggccatcccaggtgctattcatttggcgattcggcaaatgt

ttattgagcaccaactatgtgccaggccctgtgctacatgctgaagatgc

taagataaacaagataaggtcaaggacattgctcttgtagaactcttact

ttaatgggagagacagctagcccgagaaacaaacaagattatttcaggga

ttgggttaaatgctgtacagaaaatgaaactgagtaatggaatatctagg

aagcaactatatgcatgggccaactcagttactcagctttgacactgttg

ggtctagttggtacctttgtttccttttgtgttatagatggaggtgatct

ttcaaatctcctctgcagagatccaatgggataatggttgactaccattt

attgagcacctagcccatgtcaggtactctacatgcattttctcatttaa

ttccctcacctagactgtgtgctaaatggtccattagccctactttcttt

tccagtaaagaaactaaggtttggataagtgaagttgcatgccagggtcc

tgcagctttacgtggtgaatcctggatttggacctagattgggctggctc

taaagccaggcttctttcccctgctccactcttggaataaatgaaggatt

gttggttcaggatatgcagaaagctgcttagcaaaaggagagttggagaa

gcatcttggtttgagggtcttgaaggacacctactgtactcccagctgag

aggactctttctcctcatcttcaggaggatgcctccagagcctggggtta

ctgtaggcctttgccagaggcatgttcggacttagccagctgagtatccc

ctgtttcctgcttttaccatctcagctgctggtcaggaggtttgggaagg

tgacatcatctagctgaatcagctgctgcttctagatacattactggaac

tgaaagtcattaccccttccttcctgctctgcaagtctcttcagaatgag

gctggagtatgggtgttccacctgtctgcagggtcatgactggaatggta

cttgctgcttttaaagagctaaaaggtttttaaaagcttgttcatggaaa

aagctttcctcaaggcttctcaagctaagggaacagagcgtccttgagag

gcaggaggattcacaggaaggccctccccgactccctcttgtccacctac

atttagccatatgacccttctgatctcctttgcctaagtccgaggcttag

gcctagcactgtaggatgggtaaacaaaacgttatttgtggactttttga

ccttaagacttttagagtctggtcaaggggacaaaaaaaaactgcttaca

tttcagagcagaccagccaccacactattggagatatattggagatctcc

ccttgactcactaattgaaaagcaaaaaagggcaggtgtggtggctcatg

cctgtaatctgagcactttaggaggctgaggcaggcagattgcttgagct

taggagtttgagaccagcctgggcaacatggcgaaactctgtctctacaa

aaaaatacaaaaattagccgggtgtggtgtttcgcacctgtattcccagc

tactcgggaggctgagatgggagaaacacttgagtccaggaggtcaaggt

tgcagtgagccatgttcacaccactataccccagcctaggtgacaaagcg

agaccgtctccaaaaaaaaaaagaaggaaaatcgactgggtccagtagct

catgcctgtaatcccaaaactttgggagtctgaggtgggaggatcacttg

agcccaggaattcaacatcagcctgagcagtacagtgagacctccatctc

tacaaaaaatttgtcaggcatggtggcacatgccaataatcttggctact

caggaggctaaggtgggaggatcacttgagcctgggagttcatggctgca

gtgagctatgaacacatctcacagaactccaacctgggcgacagagcaaa

atcctgtctcagaaaaaagagtcattggagggttttgaaccaagaaatga

cagtccgacatacatttttaaaggatcaccaggctgggcacgatggttca

cgcctgtcatcccagcactttgggaggccgaggcaggcagatcacttgag

gtcagaagtttaagactagcctggccaacatggtgaaaccccatctctac

taaaaatacaaaaattagccaggcatgatggcgtgtgcctatcatcccag

ttactcaggaggctgaggcaggaaaactgcttgagcccgggaggtggaga

ttgcagtgagctgagattgtgccactgcactccagcctgggcaacaaagc

gagactccgtctcaaaaaaaaaaaaacagcaacagcacatgctgaggatt

ctgttcagtggtttccacttatcagtatagcacctcattaatcctcccaa

cagttttatgaggtatgctctagtggtatccacattctacagagaagaaa

actgagactttgagaggtaaagtcacctgttaggccgggcacagtggctc

acgcctgtaatcccagcactttgggaggatgaggtgggcagatcatgagg

tcaggagtttgagaccatcctggctaacacggtgaaaccccgtctctact

aaaaatacaaaaaattagccgggcgtggtggcgggcacctgcagtcccag

ctacttgggaggctaaggcaggagaatggcatgaacccgggaggtggagg

ttgcagtgagtcaagatcgtgccactgcactccagcctgggtgacagagc

gagactccgtctaaaaaaaaaaaaaagaaagtcacctgttaaagattaca

cagcaaactgccagcatccaaagcctgtcctcttccctcttccatttgaa

taggctccttccagccaacattcatgtatattctatccaggctccaaagt

ccagtaccaagccagtcttttccaggaagccttccccaagcttatctaat

ggcctgtccctctattgagagctgatctacagagctaccatccattgagc

actccatgggtgctaagtatactgctgaatgatttaaatatgttatctca

tgtaatattccaaggagccatatgccctgtgacatccattttacatgtga

aaatagtgaagctaaaatataggcacatgccatcatgcctagctaatttt

ttttttttttttttttttgagatagagtcttgctctgttgcccaggctgg

agtggcttgatctcggctcactgcagcctccgcctcctggattcaagtga

ttctcctgcctcagcctcccgagtagctgggattacagtcacccaccacc

atgcccagctaatttttgtatttttggtagagacagagtttgaccacgtt

ggccaggctcgtctcgaactcctgacctcagatgatccacccgcctcggc

ttcccaaagtgctaggattacaggtgtgagccaccacacccggccgacta

gcttatttttaaatttttagtagagacaaggtctcactgtgttgtccaga

ctggtctctaactcctgggctcaagcagtcctcccaccttgacgtcccaa

agtgctgcgattacaggcgttagccagcagtgtctggcctctaatgacac

tttattcatgagaataaatgccaaaatgtatacagtgcatgggcatatga

ctgggagaccttggtacgccagcagtggcaaggacagtcacctagaagta

agtgccctagagtacaccgaacctgggtttcaactgcccaggcttactcc

ttctcaaaaccagcccttttccagctttgtgtccctgtgtgtttgcatgg

gtgcaggcacaagtagatgacacagtggtgatgctatgcacccaggacat

ttcataatagggaaggacgttgaccgctgccatggggtaccaaaggtcat

cagggctgtgaaagccaaggacactcaggcagccttactcccctttgagt

ctgagtcatttcctgagggatattggtcattcattcatttgttcattaac

aagtatttattgcattcctgtcatatatcagggactgagctaaaggttgg

acatacagtctgaacgaaccaaagtccctattctcacccagtggacattc

caaggtgggtgacagattttaagtaaattaacatgtcagggctaggcatg

gtggctctcgcctttaattctagcactttgggaagctgaagtgggagtat

tgcttgagcccaggagttcgagaccaacctgggcaacatagtgacaaccc

acctctaaaaaataaataacatgtgaggtagtatttagtgatatagggaa

aaataaagcagagaaaagggattagggatgggaaaaggaagggtagggta

gttaggaaggcctctctgataaagtgacattggatctgaatgcagtgagg

gcatgagctgtgcggatatatgggggaatagtgttgtaggcatagagaac

agcaagccgtaaaacaaaaaaaagccagatgcagtggctcacgcctataa

tcccaacactttgggaggccaaggcaggaggatcacttgatgctgagagt

ttgagatcagccagagcgacaaagccagagtttagttgggcatagtggtg

catgtctgtagtcctagctagctactcgggacactgaggcaggaagatgg

cttaaatccaggagtttgaggctgcagtgagctatgatcctgtccctcca

ctctagtctggctgacagagcaagaccttgtttcaaaaaaattaaaaaat

aaagcaagtacaaaaaaagactctccaacatatttaaagtacagcaagtt

ggccagtatggccagagtagagtaagccagtggtaggaatcaggtctgag

acgtcaccaggggccaggtcttgtagggccacatgggaccttgctttaag

ggcagtggaaaggcatggaggatttttaagcaaagaagtgacaggtttta

gaaggatctctctgatgtggagaacaaactctatgggtaagagcgggagc

aggagaccagtgagaagccaggtgggtgataacagcagtttataccaggg

tggatgggtggtggaagtggcgagcagtggtcagagcttcaatacaattt

gaaggttggaccaactggatttgctgatgtttgctggataagaggtgaga

gagaaaggagtccaggtttttggcttaagaaactggaaagtccttactgc

catttaataaggggaagatgccttggctgatagtgatttcctgctagggc

tgtggagttgaactgcctgcattcaagtcctagtttgcttactgttttgt

ttttgttgttgttttgagatgaggtctcactatgttgcctgtgcttgtct

cgaactcctgggctcaagtgatcctcccgcctctatcttcctgggattat

aggtacgccaccaggccctgcttactgtttgactttgagcaagtgtttga

ctttgccaagccttagtttccttcaatgtaagcctactgtctagatgata

ggcagttgggtgctgtgaaaactaaatgagggtaatttgtatgaagggct

taggacaatgcctggcatatggtaaaccttcagtaagagttagctctaat

ccttattactattgttggaagacaagaagtcccagggcctcaaggtcctc

tggaaagcaggcagcaggtcacactccacctccaccaccccctcaaggag

cccgccctccaagaggcagctcagacaccatctgtccagcagagaaaagc

cacagggagacagctttcggggagcattagatagatgagcccttcccaag

aaacctaatctcattctcagcaggtgctaatcccaccaccctcccttggt

atctgagcccagcccacccacttccttccccacccccatcctgtgtccta

gaccctcttccccttctcaccagctccaagctgccaccagcaaacttcct

taccagatggaggggctctgacagaagcccaccacccatctgactgggcc

cagcctctcacatgtatcaggtcaccaagggctacaggtgccacctgtcc

agaagccccccctacaggtgccagttcctgggcgtggtgcttgatccagg

ctggagccagctggagccaggaactcccactgtgccccagctgctgcctg

ccctcccacccacaggcagagaaacaggcttaacccttcggacactgcat

atgttgagttagttgttacagttctggctcagagtattgcactgtttcag

tacactttttttttttaaagaaatttatcataaatacaacagagaatatc

taatgcatatgtacaattttaaaaataggaaaatcatcacttgtaaactc

atcacccagattaattaataatcaaacatatgtttgccttgcatgatggc

ttatgcctgtaatcccagcacttgggaggctgaggtaggaggaatgcttg

aggccagaagtttgagactagcctgggcaacatagtgagaccaccatctc

tacaaaaactaaaattaaaaaatagctgggcgtggtggtgcacgcctata

actccagctactcagaaggttgagtcggggatcacctgagcccaggagtt

taaggctgcagtaagctatgatcgcgccactgttctccagtgtgggcaac

agagggagaccttgtctctaaaaacaaacaaaaataaacgtgagtacctt

agagccccctgtgtaccccctccccaatcacagctaccccaccatccata

ctcctgagttctgtgttttcttttgtgatatcttaccatgtcaaccaggc

tagagtgcagtggcatgatcatacctcacttcagcctcaaacctctgggc

tcaagtgatcctcctgcctcagcctgccaagtagctaattttttaattat

tattattattttttttagagatgggatctcactatgttgcctgggctggt

ctagaactcctggcttcaagcgatcctcctacctcagcttctcaaagcac

tgggattacaggtgtgagccaccacacctggctacctttttttttttttt

tttttaagttttactatatatatataggtatatattcctaaacaatatag

gatattcttctacatgcttttttttgtttttgtttttgtttttttttttt

gagacggtcccactttctcgcccaggctggagtgctatggtgcgacctca

gctcactgcaacctccgcctcctgggttcaagcggttcttctgcctcagt

ctcccgagtagctggactacaggcgcacgccaccacacctgactaatttt

ttttttttaagaccgagtctcactctgtcgcccaggctgtagtgcagtgt

ggtgcgatcttggcttactgcaacctctgccacccgggttaaagtgattc

tcttgcctcagactccctagtagatgggattacaggcacctgccactgcg

cctggctaatttttgtagttttagtagagatggggtttcaccatcttggc

caggctggtcttgaactcctgacctcgtgatccaccagcctcggcctccc

aaagtgctgagattacagctgtgagccacggcgcccagccctaagttttg

tattattagtagagatggggtttcaccacattggccaggctggtctcaaa

ctcctgacctcgtgatctgcctgcctcagcctcccaaagtgctgggatta

caggcgtgagccactgcgcccagcctgtgcattcttttaaacttcattta

ggtattatgctgtgtttatcatctgcagtttccttttttcattcaaggtt

gagatccatccatgttacttgcatagcataatttgttcattttcgtatcc

atttatatgatcatgtctcaagctgtgtatccattctgctgtcagtggtt

atttgggttgtccccttgtttttgctgtttcagacaatgctgttatgaaa

actgtttgtgtctcctgcacatgcacatgttcaagagtttctcccaggca

gtggttcccaaactttagctgcccatcagaatcacctggaaggtttgtta

aacctcagattgctgggctccacccccaaagtgtttgattgtgtatgtca

gggttggggccctcagaatttgcacttctaacaacttctcagctgatgct

gccactcttggtcctgggacctcactgagaatcattgttgtagggtttac

acagaagtggaattgcttgatcagaaggtaatggcagattgttttcctgt

gtttgtgtgaatttacacagccaccaacagcatatgggttttccattact

ttacctctttgttaatacttgatatggtcagacttaaatttttatggagc

tggtgggtgtgagatggtatcttgcggttttaatttgcatttctcagatt

acaagtggggctgtgaatcttttcttacatctgttggccattcatgtttc

ctctttgtaaaatgttcaagtccttgctccatttccttttgatttgactt

tttcttatgggtttgtaggcttttaaaatgtattttgaatactaatactt

tgtctatgcatgttacaaatatagtctttcagttcttctgcctcagcctc

ttgagtagctgggattaatttgttcgccgggcgccaccacacccagcctg

tttattttattttattttattttattttattttattttattttatttttt

attttgagatagagtctcgctctgtcgcccaggctggagtacagtggcac

ggtcttggctcattgcaacctccgcctcccaggttcaaatgattctcctg

cctcagcctttcaagaagctggaattacaggcatgcactaccacacccag

ctaatttttgtatttttagtagagacggagtttcaacatgttggccaggc

tggtctccaactcctgacctcaagtgagcctccatgcctggcctccatag

ggtgttaaaggctgtaaatttgaattatttatgacttaagttatacctag

agcttattgtatatactgtattaatatttctttttttttttttttttttt

ttttttttttttttttgagacagagtctcgccctgttgcccaggctggag

tgcaatggcatgatctcggctcactgcaacctctgcctcccgggttcaag

cgattctcctgcctcagcctcccaagttagctgggattacaggcatatgt

caccatgcctggctaattttttttatatttttagtagagacggggtttca

ccatgttggccaggctggtctcaaatgcctgacctcaggtgagccaccca

cctcggcctcccaaagtgctgggattacaggcctgagccaccgtgcctgg

cctactgtattaatatctcataatgaacaaaggtaccctctttgggaaaa

agttggggctttttttccttttttcttttttaacttttcaattgattctc

aaagccagcatgtgggagagagtttatcaaaacagccaggcatcatggcg

cacgcctgtaatcccagctccttgggaagctgagacatgagaattgcttg

aacccaggagacggaggttgcagtgagctgagatcatgccattgcattcc

agcctgggcaacagagtgagactctgtctgcaaaaaaaaaaaaaatacat

actgacctgtgtgtgggatttgcttcaaaatactactggagtgagagata

gtggttggggaacagatgaaatcagattggcaccaagttggtacttgttg

aagttgaatgatgggttcgttttattctgtctacttttgtacatgattaa

atattatcataagaataagtttaaaattggccaggtgcagtgactcacac

ctgtaatcccagcagtttgggaggctgaggtggaaggatggcttgagccc

aggagttggagaccagcctggacaacatagtgagaccctgtgtctctcaa

aaaaaaaaaattagtcatttgcagtggtacacacctgtagtcctagcttc

ctagctacttgggaagctaaaataggagaatctcttcagtccgggaggtt

gaagctgcactgagtcatgattatgccactgcactccagcctggatgaga

gtgagaccctgtcacagggggaaaaaaaaagttaaaaatatccaaggact

atttccagggctacttgtgaggtagcacatcttttctttttttctttttt

gagacagagtttcgctcttgttgcccaggctggagtgcaatagcgcgatc

tccagctcaccgcaacctccgcctcctgggttcaagcaattctcctgcct

cagcctcccgagcagctgggattacaggcatgtgccaccacacccagcta

attttttgtatttttagcagagacaggatttctccatgttggtcagggtg

gtcttgaactcccaacctcaagtgatccgcccgcctcagcctcccaaagt

gctgggattacaggcgtgcgccaccgcgcctggcagaacatcttttctta

acgttttgttagccatttgtattcttcagagccaatttagtctccatcta

ctttcaggagttcctgataaagtggtcttagcaggcatggaagagactga

ttttaaagaaaacaggagagggagtggtaaggaaagactggggaggggat

tttcaggcggttgccagcccttctcctagagcctgactactactgcctct

aggattaagtttcctaccttcccttgtttcttccaggaccaccatgggat

aatttttattttctttctttctttttttttttcttttttttgtagagacg

ggatctcactatgtctcccaggctggtttcgaactcctggtctcaagcaa

tccacccgcctcagcctcctgaagtgttagtaggattacaggcataagcc

actgcacccagctgtctttcttccttccctcccctcccctgccctccccc

tccccttccctccctttcccttcccctttcttcctttccttttcttcttt

aattatagagacagggtcttgccacgttgcccaggctagtcctgaactcc

tggcctcaagcagtcctcctgcctcggcttcccaaagtgctgagattaca

ggcatgagccactgtgtccagccaatttttattttaatggtattcttctt

tccctattattttacctggacagacttaacccagaggatagaggaacctc

atcatggatcctggtccgcttttagcttaacagcatccaaagacaaaggg

tcagagatggtagcaaccctagtgtggttgattacacagtagtggtgtta

ttttgttgtcataggatgcaaatatcaaagttgtatctcttaagatcttt

ctgtgtcagtacataaaagctttctcagtctttttaatgctgcatagtat

tctacacagggcactattctatactttaaccagttcccattggtagacac

atttttttccagtcttttgcagttataatagtgacactcaatgaatagcc

ctgtatatttgtcatttggcccatatatgaatacagtatatctggggggg

atgaatccctaaaagtggaattactggggccaagggtatgtgtacttggg

attttaatagggttagttagcctgggcccacagcagtgggtaccagttta

caggagaatcttttttcttaacacccttccacccaagtggagagtttggt

tcagcacatttctttttctttttctttcttttcttttcttttctttttct

tttctttcttttttttttttttttggagacagagtcttgctctgtcaccc

aggctggaatgcagtggcacgatcttggctcactgcaacctccacctcct

gggtttaagcagttcttctgcctcagcctcccaagtagctgggactacag

gtgcccaccaccttgctgggctaatttttgtatttttagtagagacaggg

ttttgctatgttcgccaggctggtctcaaactccagacttcaggtgatcc

acccacctcagcctatcaaattgctgggatttacaggtgtgagccaccgc

gccggccatcagcacatttcttgagtgccttcctgtacctggccctggat

ataagataaacactttcatgtatcaagtgaaattgttactatttgccagg

caccagcacattatctgataatcctcataactactcaaagaggaaggctt

ttagatgagataagtgaagctcagaagggttaaatagggtgccatggtca

tggtccagctattaagtggcagagctaggattaagcccaggtttcaatcc

tctgatgcctgtgctcttacagtacacctttacctgcctctcaggaggaa

gaacatatggtcttattgtctaatacagataaggaatcaggctcagagag

ggtaactggccacaaaacagccagcctggaagctatgaggaacccacata

tgtctgatcagaggtctctggtctacactgagctgcctgcctccctcccg

gggaaagagtttgagaattttcttttttttttttttttttttgagacaga

gtctctttctgtcgccaagctggaggagtggagtggcaatcttggctcac

tgcaacctccgactcccgggttcaagcgattctcctgcctcagcctcctg

agtagctgggattacaggcgcgtgccaccacacccggctatttttttttg

tatttttagtagagaaggggtttcaccgtgtttgccagaatggtctcgat

ctcctgacctcgttatcctcctgcgttatcctccttcctttgggaggccg

acgcctcccaaagtgctgggattacaggcgtgagccaccgcgcccgaccg

agaattttctccgttcagagcttcacttgctatagagagcctcaagtcac

tggttctcagtgtgcatctattggccatcagagggtcctttagggatgac

ccatctgtgcttaatttactaatgacagtatagttgctgctatttattga

gtgcctgccatgtgcaagatgctgtaaggtacagtgtagtttgttttgtt

ttgttttgtttttgtagagatggaggtcttgttgtcttgaccaggttggt

ctcaaactcctgacttcatatgatcctccctcctggacctcccaaagttc

tgggattacaggtgtgagccaccagactcggccagtgtagtatttttaat

cttcacaataatccttcttgggaaacatgttattgtcctcattgtccaga

ttagaaaactgagtgtaaagtaagttaaattatagtcctaaggttgaatg

ctaataaagacagaatacaagtccaatatattggactcaaaagccctcac

ttaactatggtctccatgggcttcccttggctctctctgcctttttttat

tttttcttattgcttgaggccctttctggaaggtaagtctggattatcta

cttcacactgttttagagaagacttgtggtttccatttaccccttactcc

ctccgctccatggcctttcagggagaacactgtgggtatcatgctgggtg

gcctggagggtccaagtaacaggaatctagaaggatggaccagatgtgaa

caaaagaaagcctgagtaggacacaaaacagagaagtggggctgtaacat

ctctaagatattacagcttgctacttccactctctttgcaaatgtggtga

aacccaggctggagtcataaaataatagcataggatcattaactaaagtt

tgtctagtgcttccttgtgttcacacattatctcattgaacctctgacga

tgctaggaggaggtaaatagggttttcctcttaccttgggtgaactgagg

tcttctgactaagtctcaggtcctttctaccatttgtgctgcatccacat

ctctactgggaagaacacaggctctgcagtcagacacacctgggtttgaa

ccacctccattttgtgtgtgttggatcttgagcaagtgaattcacctttg

tgaccatccatgtcttcataaaataagaattttaaacacacatgagggcc

tggcatatacaaaacattgaataaatagcagtgatctttttattgacatc

agaagggggttggggacccagaggctatttgggctcctatggcatagctc

ttggaacgtgttaattttcaaggacaggggtttgcaagtcagagcaatgt

caagtaaagaacaggttctttgggaggccaggatatgctaatgggtgaac

aatgtaagggaggcataataataatgccaaaggcctcctagagtacctgt

tcactgcacttttgggtctttggtttatatatgaagaaaatgaggtgccc

ccagaggggaagtgggttgccattgttacacagtaagttaataaatggca

ttgctgagattagaactcagtcttttgtctgccaagaaagcttacttttc

ccttacactccactgctcccttgaggggaaaaaaaaagggcccggagata

ggatcttagtgctttggaactggagagaaagccttagagatcatctgatc

catcctccacacttttcagatgagaaaagtgaggcccgtggtgcttaatt

gacaggaccatagccaaatagagccaccacaaagcaggagtaagctcacg

tttcttgaacctgaccatgtggaactttcaaattgccttttttctgtgcg

gaactagagcctcactgaaaataccagtggtttacaataagaacagctgt

ttagtttggggaacatttttccaaatagaatcttctttttctctctttct

caattctttttttttttttgagatggagtcttgctctgtcactcaggctg

gagtgcagtggcacgatctcagctcactgcaagctccgcctcctgggttc

acgccattctcctgcctaagcctcccgagtaggtgggactacagacaccc

gccaccacgcctggctaattttttgtatttttagtagacggggtttcacc

gtgttagccaggatggtcttgatctcctgacttcgtgatccacccacctc

ggcttcccaaagtgctgggattacaggcgtgagccaccacgcctggctgt

ctttctcaattcttaataaaaatatactagaagaagaaaaagctttaatg

gaggaccttggtagaaagcaggtaaaaattatatagagtctctaaaatct

agggaggtccccagagccagtaaatttagaggccagccttctcgcctatc

ccagcaagcttatcacttgtcttgccctattcacaacaggccttcagaag

gcccagccttcccttggcagctgcatgtggtgaacaccaacggaaatccc

tggtatttttctttgtatgcaaccaaccgtggagatgtaggtcaggcatt

gctggcttggcagttacaacatctctagcttcacccctgcacctgtagcc

gtgttcactaagcccctcaaagatgggaaatggatggtcccttcttaata

gggtttggatccccgagggctcatatgggtggagtgggtttgggaggaga

cagaccatccggaatgtggagatgagaagggtagatctggctcttccctc

cttaatgtctgatgtctggtgttactatagctggattggtggtgcttctg

ctcagctcaccttagagtaatcatcactgacggaggagactggggaggtc

ccatccttcccctgggctcagctgggtccccagacctacacctgaacccc

acgcctgtctggtgtttacggattgaagctgccaccctttgggacctatc

ctgttttcatttctgctctcagctctctggtctctgtggacctcacttac

aggtttgtacgtgcctgtcacccacggtgcagccccatgaggctctccca

gcctggcacactcatgcctctttggcctttgttctcttcaggaaaactgc

tagctctatgccttgctagggagggcttttctggcccctcagataggttc

ttttggtccaacctagctatacctttcctgagattcagtctctttgtcca

caccctgggggacagggctgtggttccaaggcatgttggggggtacctgt

gggccattctggactggcttccggtcagctggctgactctgccctgggtc

tcaaatatctcctctgtaaaggtgttagaccaggatcttcaaggtaactt

agaggtgatatgcagatggacattttctgttttaatagtcagtgtatttt

atgttgatttccatctggggcaaatgatactctttataggaagaataata

agttgccttttcattgcatttatttatttagcaaatatttgctaagtgtc

tactatctgtcaaccactattctagctactggggatacctgccttcatga

agttcacattcttttttttttttttttttttttttttttttttttttttg

agatggagtttcactcttgttgcccaggctggagtgcagtggcgcagcct

cggctcactgcaacctctgactcccgggttcaagcgattcttttgcctca

gcctcttgagtagctgggattacaggtgcccgccaccacacccagctaat

ttttgtatttttagtagagacggggtttcactgtgttggccaggctggtc

ttgaactcctgatctcaggtgatccacctgcctcgttctcccagagtgct

gggattacaggcgtgagccactgcgccgggccatcatgaagtttacattc

taagaggacagagagtgattaggtggccggtggtaggcagcttttggtat

gtgtttgcctttttaataaaggataattaaggaaggcctctgaggagcag

gcatatgagcagagatctgcatgatctgtctgagggaggaaaccatgtaa

cagtctatggaaaaaacattccagacataaggaacaagtgcagaggccaa

agggtgggaatggtgtgcctggcatgtttgaggaagagcacgggagatca

gtgtggctagaacagagtaagcagaggagaagtgattggaaaggaaagtt

tgtgtgagcagtggtgagacattttggttcttcctctgggtgagatggga

aggcagtggagggttttgagcagacaggaaactgactttgctgacccctc

tggctgccatgtggtgaaggaaccatggagtcgataagaagctgggagac

cagtaaggaggctgttaggacaatacaggcaagaaatgatactggcttgg

accaaataatagcagtggaggtggtgggaaatggtgggactctagataaa

tttggaatacagattcatcaggatatattgacagatgagtgttagagggg

atttaaggatgagaaagatgcagaaagaacaggtttagggaataaattga

gaattttcagtacatgtatttaaatttataaaaagtaagtaaattgatag

aataatgttaagtaggccaggcacagtggctcacagctataatcccagca

ctttggggggccaaggcgagagaatagcttgagctcagtagttcaggacc

agactgggcaacatagtgagaccttatctctacaaaaaaagaaaaagaaa

aaaaaattagccaggcatggtgatgcacacctgtagtcccacctagtcgg

gaggctgaggcaggaggattgcttgagcctgggaggttgaggcctcagtg

agccttgatcacacgattgcactctagcctgggcaacagagtgagaccct

gtcttgaaaagaaaaatgttaagtaaataataacagaatcacatataagg

tcaaagcagtgatggctgaaattgtggatagagtggaccaaatgtgtctg

tgctgccttccagctctaaaatcctgcttcagtgattcagaccacatagt

ctcccattcacagagaacaagaacttgaagctcagcctagaataagcaga

taaagtgacaggagagcaatcctttggaccaggcctacctcctgttaatt

tggaaggacttgggaagttcttgcacatatactggccttctggcattacc

ttatttatttattcattcattcactcagtcagtcagtcagtcactgagca

ccttctgcctagtaggccctggacaacggcagctctgatgcttcttctct

cctaatataaagcaagctaccctgacccccgctcccactggagcctgtct

actctgtggcctggcatatgcctggcggtaatgtgtgtgctgaaagtgga

atagccacagcctaggcagagccctgccctccctttgctgcactgtctct

gtccatctctgccttcctcatcttattgccatccacactttgaggcctgg

tgagcaaatgattttgcttccatgagagcagcagaaatagttgctttatc

tctttcttcatataggtcctcatagttatcaaactggaaagaaatcccca

agggcctcttctctcgccttttatagatgggaaacagagggccaagaggg

caggcaactcccacagtccacatctaattcatcagcaaatcctttcagct

ctacttttattgattgattgattgattaagaacagtatcttgctgtgtct

tccaggctgaagtgcagtggcgcagtcatggctcactgcagcctcagcct

cctgggctccagcagtcctcccacctcagcctccggaatagctggaacat

gccatcacgcctggctactttgtgtcatttttttgtaaaaataggatttt

cccatgttgcccagactggtctgaaacttttggactcaagcagtcctctc

acctcggcctcccaagtgctgggattgtagacatgagccgctacacctgg

cccagccttactttaaaaaataaattctgaaactgactgcttctcaccaa

cttccctggcaccattgtggctaagccactgtcatattttcctgattatc

tcaccagacccctggagatctccacccagcagccagacggacgctattta

acaccaaggtcaggtcatgtccttcccctgctcaaaaccctacagtgctt

tcccaggacaagcccaagtctttaaatggcctaccaggccccacacaatc

gggcccttggctacctccagatctcatcaccaccagtctcctcacattca

ctggacttcagccacactggcatttttcctgttcctggagcacctcaagc

gtgttcctgcctctgggcctttgcacttgttcccacagccttttctcaga

tctttgcagtgcctgtttcctcatttcattcaggtctctgatcaaattca

cctccttaagaagttgtccctgaccatggattctaatcttaccttcttcc

atgtttctctgtaccccctttctcacatttattctttcttaatgcatcac

cgtctgacatgtttattgcctgtctcttcccatttgattataaactcata

caggtacgaattttcattctatttagtcactgctttatccccagcgctta

gtacagtgcctgactcttagtgggcacttaatgaagatttgttgaatgac

tgagtgtccaaagctagggaaggcagcctgtctgggccttcctacaactc

cataggtcagcctggcttacaaataggtcatgtcccatcttagaaaactc

atctaatagcttcaccaaagctacgtggcatcacttttgtttccaagata

aagatcagacagagggaggaactctctctgcaagtactgtccttgcagat

ctggcttaggggaaagtgggttcatcactgagtagcctcagcacactctt

ccctttctcacccctcaaggcacccagggaaccaggggggaatgctactt

ccatgtacttaaagtcatctctgacttcgaaggcctaagaatagcagatt

ctcagtaaatattgggtaggtgaaagggtgaaggaaagaaggtcttagta

aaatctcttcatgtatcctgttgcactggtaaatttaacatgctgtgaag

ttgaatgccctcactagtaaccaggggcctcaggcaaagtctcatcactt

cgctgaattttagtttcctcattggtaaaatgggaacagcacctacttgc

aagattttgagaattcaaacagtaaatattagttcagcaaatactagctg

ttactttttgtttgtttatagctacaaataagagagacactatgagaagc

acaaatggagccactgattttttagctataaaaaaacaggctgggtgcag

tggctcatgcctataatcccagcactttgggaggctgaggcaggcagatc

acttgaggccaggagtacgagaccagcctggccaacatagcgaaacccca

tttccactaaaaatacaaaaattagcctgcaaaaatacaaaaattagtgt

ggtggcacatgcctataatcccagctactcaggaggctgaggcacaagaa

tcgcttgaacccaggaggcagaggttgcagtgagcagagatcacaccact

gcactccagcctgggcgacagagacagagtgagacttcgtctcaattaat

aacgataataataacaacagcagcagcagcagctaacatttacagaacgc

ttaccatggaccattgaactctgctgctaggcctgtgaagtaagtacagc

tgttttcattttgcagccggaacaccaggcttggagggggaaagtgtctt

gctccaggtcctgcaaccaggtagtcacagaactagggttggaagccaca

ctgtctcactccagggtagggaggcctgagaccaggctgtgctctgtccc

tgtatgaagccttggaggtatcccttcacttcctctgggcctcagtttct

tacctgctcaggatcgaaccacacatctctgtgacgctctgaaatttctg

gtggtccacaggccacatcaagtgggacagtgagttgcttccaggcctcc

tggtggtggtgctctgggtattgactgtgaacagcacacctctgggtgcc

ccctccctcgacatcaggcaaagtaccctagccctcttctcaccatctca

ggcatcttactgttctgcaccttaaaggggtaatggactggggcagaaat

gaatagggtagaaaacaagaaggaacccactcccttgtctggtgcagatc

aatgttggatgctgcagagaagagtatgaggctgacagtcagtggatgtg

ggttttgtttcccagttccatcacatcacagaatagaaggaaatgcaaca

ccgtgggtgccttcagccctgagtttttggttacatggggtgtgagcatt

catctgtacagtggggagagaagggtgttgcagggatttgctcccttccc

caaatcctccagagctgttgccttctgctgtggaattccaaacatttctg

atcagataaaggtggctgaaaatacttgggacttgtctggactagttgga

ggcggtgctggggtgaagaggtcgaggccacaggagctagagcgtggaaa

gatctagccttgcccaggagtctgcctggtcatagatgtgcctattgctc

tcttcccctggaatcctgcaggttcagagcaatgcccagcgggctttgag

gtggagaaggatcagactcctttgtggcaacaagttctcctacttcagaa

tgttgcttctcaatttacacactaattactggatcttttggatacccctg

agagagagaaccctggtagtcactcatctgtctgcatgagttactttcat

tctccttatgctgtagccctagacggagccctttgagatgattcccactt

catttgacagttggggaaaggcccatactttcccagagtctcaggcaggg

gaaaggactcacctgcaaatcagcgatggaactgggaaacaaaatccaca

tccactgactgtcagcccagtactctcttccccacagcatccagcactga

tctgcatcagacaagggagtcgccatagctttgggtggagaggaggaagg

agaagctgctttggactatggagcacttggggtcaaagacctcagaaaga

agtcctagtttttggagattgctaccccattcctgcctgttgcctcagca

cagtgcaaaagatgccttttctagaactttcccataagcttcctgtcccc

tttccttatcctgagagaaagtgtctggagaatactggacaccttgcata

atcacctgctatattggggcctctttctagttttgccaggcagtgaggaa

agggatgaagagagtgggaactggcctttcttctcttccctcctctctgt

ccaacactcagccagactcttcagcaccttctctggacccagcactgggc

aaggcctaggggacagagcccatggtcagatgggggaagaagtgagcagc

taggctccttgagaaagtgacctgtggattaagacctcagagataagtgg

gacagagcttggtagaagcaccagtgtgggcaaaggtcctgagtctgaac

agaacatggcatgtgaggaatgaagcagcctggccctaggtgaagctgag

aaaaccctgcaggcccttgtaagccatagagaaggctccgagacacaagc

tgagccatggtctactagtaagaacaataaaccagacatttactcaaagt

taactatgtgccatgcacaattctaagacctttacatgtattaatttgaa

tcctcatggtattcttttgaaataaggacattttgggccagatgcagtgg

ctgacgcctgtaatctcagcactttggggggccgaggtgggtggatcatt

tgaggtcaggagtttgagaccagcctgaccaacatggtgaaatccagtct

ctactaaaaatacaaaaaaattagccaggcatggtggtgcatgcctgtag

tcccaattacttgggaggctgaggcagaagaatcacttgaacgtgggagg

cagaggttgcagtgagccaagattgtgccactgcactccagcctgggcga

cagagtgagactctgtctcaaaaaaaaaaaaaaaaattaaataaggattt

ttttaatccgcattttacagatgaggtcaaaagcaaaatatcttaatcta

gtttgcaagacttattggttctctgcctgctctgtgctctggcagcccag

cttggtagagaagtagaaattaattgcctttatccagatagcacttccat

ttcacaaatgagacaactgatacttggaggagagcaggagagcagcagcg

gagagagagcagaactaagaccagccagtcaagggagacagagggagacg

agcatcagctccctgacagggaagagttttccaacggggctgtccgaaga

tgaaatgtcccacccaggaaggagggagctctccataagtgcataaggag

agaatgccctcgagatgactaccaactcccatgggtgtggaccatcagaa

tgaaacagacccactgtgggacttcagcactccacccttctgaaccctgt

ctgtgaaatgggggtgttcgtggtgagcttgagtattacatggaaggttg

gatgggcatgctgtaaggtccctggagtctccgactctgttccaagggtg

agtattggtttttctcaccctggacactgggagaagaatggagggctgtg

gccatctacttacgaaagaatagggagtcaccagggttgaaatgagagag

ttgagatgacgggaggcatcggaatcccaggggatgagcctatatgagca

catagcagccacccaggaacagttaaatgtttggactctggagacagact

gcctgtattcaaatcccaactctaccacttgtggcctctgtgaccttggg

agggtgatttatcctctctgcctctgtttcctcatctgtaagctggagag

taataacagcatgtactcggtgggattattgtgaggattaagcaaattct

ttacacacgaagcacttagcaccatgcctggtacttagtaaacatcagct

gttttatctccatcggttccagggaaggaggttatgagcccccaaatgag

catctctgcccccagtaaatgcttttctgaattctctctctttgccccca

agcccatcaccctggttctgagccagagagatggagagcccacactcgca

ctgctaggcctgatagctttctcgattccccagcggaggtgttatctcct

cccggcacaaccgcttccttttctgagaaatgcaagtgcatctgtcaggc

acagcttcttagaactaatggtgggggaggaggtggaggaggagaggctg

gcagctgaattgaacaggtcagagttggaggaagataaacatgaagtttg

ttgtgaggataccaccaaatattccctatagcccagatggcttgagccct

ctatgacaccccctgatctcaccactactgctagggaggtcaaggcagga

gaaactgaggctcagaagccaaaaggcttgctgtggcacccacccatacc

tgggccggctggactgaggtgttgaccccaacacagtcttgccacccctc

tctttgccctgcctccactgagctgcttctccaagttgcccacctccagg

cctatgtttccgtgggggagaggagaaaggccatgttcttctcacatctc

gggagccaacagctcccctgggaaaaccagaggaagtggattccaaccac

aaacctcacttcttccgagtctccgcctgcaactccctgccaaaagagtg

ataaggcctctccgtttgccctacccccttcctaccatctcccagcgatc

ttctagcaaggccatgtttgtctattcagcacacatttatcagcctctcc

ctcagggaaactgaggccaggagcataaggtgactctgcaggtgactcac

agcctgccaatgcagggatgtgagaaggacagaacaaggtacagttctgg

tgtccctggcacttttcctggctgctagtagcctcttcctcttctccacc

tacctctttcctgaaataaccaagcaggaaggaagcaatgggcctgggcc

ctccagagtcacccaagaggaaggaagggttcgtctggcacaggcatgct

tgttcctcactgaagccaggacggaagggtttcactttccctcctggtct

tcgtcacatccttttcctgtggcaccagttcccctcagctggcaggagcc

agtgcttattcattcagctgatggcctcaggctactttaccctccttgac

agacccagggccatggccctggctcagaatatggggtttgctgccaaaca

tactggggcttaaaaatggcttccctttttttttttttttttttgagacg

gggtctcattctgtcgccctggctggagtgcagtggtgctgtcacagctc

actgcagtttcaacctcccagcctcaagcaattctcccacttcagtctcc

tgagtagctgggactataggtgcatgccaacacactcagctaaatttcta

agatttttcttatagagatgggggtcccaccatattgccctggctggtct

cgaactcctgagctcaaacagtctgcccaccttgaactctcaaagatgct

gggattacaggtgtgagccatcacagctagctgccacttatccactttag

ctaggtctcagtatctctgagcctcaggttacctggacttagaatgaagg

ccttcctcagccacagaagtacatggacctggcccatgattgtcctttag

aaactgagcagagtactgcatcaacgttaggagtaaaggcactgaaccag

attgctggtgtaggcatcccagttaggtcaccagctcactacatggcctt

gagcaattcacagatcatctgtgggccacatatccatcagctatatgggt

taatggtagtacccactgacagagtgtagagacaactaatgaactaatct

acaaaaaaaccaagaagagtaccttacagagtatgtgctcaggaagtgtg

cgctgctgctgaagtcactaaggaagtcaggttttctgtgggtagagata

gcagggtctaagatggcaagattggacatcagaatgagtgtgaatgaagt

actatcttggatagattaaaagaaaaagaatggtgtgaggccagaactca

gacagatcctactcttgccacttcctagatagggatcttgggtaaattct

ggagcctccattttcacatatgtaaaatggggctattgataaatgttatg

aggatcaaatgaaagcattcatttgttcttctaaaatagatttatttaga

atggcatctgctgtgttccaagtatagcacaccaggttctagagatacag

gtgacagagccacacacagtcaagtgggggagacaggacaaaaaataatc

tctcaagtgaatatgatgattgtacatgtaattgggaaatatggcctagt

gtgcatagactgctttatagctcagtgcccagcacaggtgaaacactgaa

atgtttggtaatggtcagctgtcataataatagttttttgtttctttgct

tttatcttttgtttttttgagacagggtctcactctgttgcccaggctgg

agtacagtggccccaaacatgattcaatgcagccctgacctcccaggctc

aaagatcctcctcagcatcctgagttgctgagaccatagatgcataccac

cacacccagctaattttttttttttttttaatttttttgtagagacaggg

tcctcctgtgttgcccaggctggtctcagactcctgggctcaagcaatcc

tcctgcctcagctcctaaagtattttattactgttaacaagtaagggctg

ggtgcggtggctcatgtctttaatcctagcactttgggaggccaaggaag

gtggatcacctgaggtcaggagttccagaccagcctggccaacatagtgg

aaccccacctctactaaaattacaaaaaattagccaggtgtggtggcggg

tgcctgtaatcccaggcaacttgggagactaaggcagaagaattgcttga

acctgggaggctgaggttgcagtgagccaagatcatgccactgcactcca

gcctgggcaacagagtgagaggctatctcaaaaaaaaaaaaaaaaaaaaa

gcaaggacaccatggaccagagaggtgaagtaacctgctcaaagtcacac

tacttggaagaagtgggaagggtgggcacagttgctcatgcctataatcc

cagtactttggaagaccaaggtaggaggatgaggccaggagtttgaaacc

accctgggcaatgtagtgagactccatctctccaaaaatttacaagaaaa

aaaaaaagaaggaaatgtgggcagtgaagccaggatgggaactaaaagat

cagattcaagtagggactatacttcagcctctggctggtcctctggcact

tgggcccctttcctcagggcatcatgccaaccttttcacctgacttctac

agggtcacaggcacctacaaggtgacctgcatcgttcagttggctgcctc

tgggcagagatctagctgaagtcagtctgtgcctgtctgagtaggtcctg

ggccagacttggtccctgcctgcactgagcccccagactgttgggggaaa

ccaggcacacagaggccatcaacccaggaacacccctcctctccctccca

cctcagggcctttgcacttattacctctgctaaaacaggctttttttttt

ttttgagacagggttttgctcttgttgcccagactggagtacaataatgt

gatcttggctcactgcaacctccatctcctgggttcaagcagtcctcctg

cctcagcctcccaggtagctgggattacagatgcctgccaccacacccag

ctaattttgtatttttggtagagacagggtttcaccatgttggtcaggct

ggtcttgaactcctgacctcaggtgatctgcctgcctcggcctcccaaag

tgctgggattacaggtatgagtcattgtgcccggctcccccagatattct

ctgtcacctccccaggacttaaaaccctatcatctctaaaattatctttt

ttttttcttgttatcctctctgcttcccagtaagacagggtttctcaacc

ttttttttttatcatccacccctgcaaggagtctttttcagatttttttg

tcctaattgccctcccatgaaatttgaatacctcagctatactatactat

atatagtatatatagatatagatatgtataatatataatatatagtatat

gtaaagtatatatatagtatatacagtactatatatactacatattgtgt

gtatatatatggtatatatagtatagtatagctgaggtattcaaatttca

tgggagggcaatacctatatactatatatactatatactctatatctcct

tatgtgctgtggccctgtggagagccacaaatcactgtaatgtctaagaa

tgcttcacttcacgccattggaggtgatcttaccaccattgagaatggac

tagaacataagctccatggaagccttgctcacggctgagtctccaatgcc

agcctcagtgcatacccctggtataaacagaagtccaaggagtctgggaa

cagaaaggagacattcatcccagaggttggggagggatgagcagggaaga

ggggactccaaaatggagtcctataggcttagtaggggttggttaactga

ggaagatatggagtgaccagttaaaggacagtgttggcaaaggcaccagg

gcctgaagcactcaggagtatgtagtgtggggccattccaagtgggtggc

caggactgttgaagcctcttgaagtattggaagaaggaccttgaaagtga

tactagcctgggcatctcctaggggccaagtcaggagacccagttttcta

gtgtaggctctactacaaaccagcagtggaaacgtagaatgggcaggtgg

gttggccttgctgacttcatgggcctacctggctaggaatccatgactct

cctgtggaggtcattagccattgcctgtcgccagtggccctacaaacccc

cacatggaagtcagctaccccactttcctcttaactcaagcagaggatgc

agacaggcccttgggcctgggtctcctagaaacctttgtgacatcagaga

cctccctacaacccactcttaaacctcatcccctgcctaccatgccagcc

tggttaaaaccagtgggagccacccaaggtgcagtaggggcttatctctg

cttttccagaccagggctgctctgcaatcctgtttatttatccagagcag

ccgcctgctgagaaaatggctctgtttatttccaagtgtgtcgtgtgaac

catgggctaaatatacccacgccaccgcctctccagcacccgccctggaa

cagcacgccccattttgctcatgctgtctccgcaaaagcaatccttagaa

tccacaccccttggaaatactactccccttccccaggaaaagaagctgaa

atggaaatgaaggttgtcatgaccaggaacacatgagtggaagctggggc

aggtgaaatgagaatgtttctatggcactgaaatgagaatgttccctcca

cccaggaatgactgtttctggacagtgaatgctttgtgcccttgtgggga

cagcgcctgtttactccaacccagccccagagctgaaaatttcctgtggc

tgccagccgtctacttcaggaaagttcccatcaagggagccaggcctctc

ccattttggctgcaaccccagtgtccctggtttcttgagagacatttggc

ctcttgggataaccagggccttctcctatgctcttcatagtgtggaactc

tcccccatcacagataagcctattttgtgactccacctgggtgtacttgg

gagcaacaaaatcaaacctcctgtaagaatatttttgtctgccaagaccc

gcagaggctagctaggatggagagggagacagggaagcatagccacggcc

ctagtgccctcctttccccagatcctccacctcaaccctgctcccacatt

ccttctcctggaaatccatcctttcagcactactgacaggacaggtccct

ccagcctccaacttcagggaagtgtcctgacatagcagccctccctttga

agctaggcagtcttgcagtttgatcccgactctgctatctgtttgctgag

tgttccctgctttaaatatcaggcatcttatttgtgaaatgtggctcaaa

tctacatctcaggattctgatgaagatgaaatgggataccaatgcctgac

atgtagtagacatgcaatacatttgttaactcatgtattctcatgtattt

acagagcaagacactgttctgagcactgggtgtataggaacgagacagga

cctctgccctcatcaagacacatctagtataggccgagggcagtggctca

cgcctataatcccagcactttggggctgaagagggcagatcacttgaggt

caggagtttgagaccagcctggttaacatggtgaaaccccatctctacta

aaaatacaaaaattagctaggcgtggtggcaggcgctactcggggggctg

aggcaggagaatggcttgaacccgggaggcggaggttgcagtgagccaag

attgcaccactgcactccagcctaggtgacagagcaagtcttcgtctaaa

aaaaaaaaaagacacatctagtatggaagacacatcacaacaaaaacaaa

gtacagaatctagttatgagaagtacagcttaatgggggaaaaatgtggt

gaggtggtgaggccgggcatggtggctcacgcctgtaatctcagcacttg

ggaggctgaggcagaaggatcccttgagcccaggagtttgagaccagcct

aggcaacaaagtgagaccctgtctctacaaaaaatttagaaatattagcc

aggcatggtggtgtgtgcctgtagtgtggcagctactcaggaggctgggg

caggaggatcatctgagcccaggaggtcaaggctgcagtgagccatgatt

gcgccactacagtctaacctgagtaacagagtgagacactgtcacaaaaa

tttaaaaattaaaaaataaagcactgagaagaaaagctggcatggtggtc

attctgtgtagtgtagttcctgtaatcaggaggctgaggcaggaagattg

cttgagtccaggagttcaaagtcagtctgggcacatcttttaaaaacaag

aaaaaaaaaatggtggctgggcatggtggctcatgcctgtaatcccagca

ctttgggaggccgaggtgagcagatcacctgaggtcaggagtttgaaacc

agcctgacccaacgtggagaaaccccgtctctactaaaaatacaaaatta

gccaggtgtggtggcacgcgcctgtaatcccagctacttgggaggctgag

gcaggagaatcgcttgaacctgggaggcggaggttgcagtgagctgagat

cgcgccattgtactccagcctgggcaacaagggtaaaactctgtctccaa

aaaaaaaaaaaaaaaggccagatatggtggctcgtgcctgtaatcccagc

actttgggaggccgtgggcggatcacttgaggtcaggagtttgagaccag

cctggccaatatggcaaaaccttatctctactaaaaatataaaaattagc

tgggcatggtggcacatgtctgtaatcccagctactcaggagaatcgctt

gtacccaggaagcaaaggttgctgtgagcccagtttgtgccactgcactc

cagcctgggggacagaataaaactgtgtctcaaaaaaaaaaaaaactgag

aagaaaaatcaaggaaggtaaggggacaggcagagagagacgggggtggt

gtactgttttagacaaggcagtcagggaaggcctcccctaggaggtgaca

tttgaggggagtcatggttgtgaaatgagggagtgaggcaggcgacgacc

tggggtcaggtgctccaggcaagcaggaacagcagactcctgcggcagga

gacgttggaggaacagcaaggaggccagtgggactggagtgaagttcagg

ggacaggggtggggaggaggccagaaagtgttggagccagatcccactag

accttgctggccagagtaaggagtggggactttatttcaggtgtgatgga

agccctgggaaggttgagaacttggagtgactgaatggagtgtgcttctg

caagggctgagattcatgggtcactgctctctactgccttaatgttgagg

attccttcagtcaaattagcagctcagtgcacatggcattttgctaggta

ccagggatacaaaaataaatgatgctgcccttctaagtgcaggagaatgc

tgctggagggcttccagtgttcttagaaagctctgtgtggagcatggact

gagggggttgtcaagactggaagtgggccacggcaggggtcacaagtgag

caaattgataggtgagacctgaggccaactggagatggcatgcctgtctt

gagggagcagccactcggccatttgggactggttggaaatgtaggcccag

tgtggcagaggtcttggtagagaggcaagaaccagccagtaactcattca

ttttacacacacacacacacacacacacacacacggtactatggtacttc

cttttttttttttgaacggagtctctgtcacccagactggagtgctgtgg

tgcgatttctgctcactgcaacctccgtctcccgggttcaaacaattctc

gtgcctcagcctcctgagtaactgggattacaggtgcgtgccaccacacc

cggctaatttttgtatttttagtagagacagggtttcaccatgttggcca

ggctggtctcgaactcctgacctcaggcaatccacccacctcagcctccc

aaagtgctgggattataggcatgagccaccgcacccagcctggaacttac

ttctttcttacatatttctaggtgccagatgccagttttagattctggct

ccaccactgacacagctgggtgaccttgatccagtccctggaccctcaga

ggcacagcttcttggcctgtaaatggggagagacagcctgcattgcagaa

ttgttgggaggattaaaggacaaattgggtactacgcttactcataaaag

ttatgttctaagcgtccttctccctgccccacaggctgctgcaccctggg

ctgggtaatttagcccacagctcctccttccccctgtcatgtgcttccct

tctgacttagattcagggcagagggctgttgatcacaacctcctgggccc

agagtcacaccccctgcagtggaggatgtccgggacaaggcctcacagac

cccgtgatggaggcctgtcagtcctgcatcattgttaccctctctcctct

gccctgctcttcttgtcaccaacctgcccaccagaggctccctccagagg

gaggagcctctggctgaagtcccaggacccattctacccctgaattggcc

tcaaatgaccctggggatttagccctgggcttccttatccctccctctgg

tgcagcagtagaaaatctgaggcctagtggctccacactccacactaccc

tcagactcaactctaggctccttgcggcttccaggccacgtatggtctgg

ctcttgctcacccctgcaccatcatctcccccatgccctcctctcctgca

ctctgctccagccacctggcttccatcagttccctgaccctgcccatctc

gttcccaccttggattctggagtaagtgctacttcttgacttacctcaga

cagggcactttcctcctctctacctgtttctcagcagtaacatggtttga

ttatcaccaatggagggtcaagaaagaggaccacccttagccctttacct

ggcccagagtgggcacagatggctacagtttttcacatcaccaaggccat

agcatgcagatggcccctttccccacccacctacatcactctctgaggac

taaaactgcaccttccttctctctccctgcag	GATACTTTTGTGGAACTC

TATGGGAACAATGCAGCAGCCGAGAGCCGAAAGGGCCAGGAACGCTTCAA

CCGCTGGTTCCTGACGGGCATGACTGTGGCCGGCGTGGTTCTGCTGGGCT

CACTCTTCAGTCGGAAATGACCAGACACTGACCATCCACTCTACCCTCCC

ACCCCCTTCTCTGCTCCACCACATCCTCCGTCCAGCCGCCATTGCCACCA

GGAGAACCACTACATGCAGCCCATGCCCACCTGCCCATCACAGGGTTGGG

CCCAGATCTGGTCCCTTGCAGCTAGTTTTCTAGAATTTATCACACTTCTG

TGAGACCCCCACACCTCAGTTCCCTTGGCCTCAGAATTCACAAAATTTCC

ACAAAATCTGTCCAAAGGAGGCTGGCAGGTATGGAAGGGTTTGTGGCTGG

GGGCAGGAGGGCCCTACCTGATTGGTGCAACCCTTACCCCTTAGCCTCCC

TGAAAATGTTTTTCTGCCAGGGAGCTTGAAAGTTTTCAGAACCTCTTCCC

CAGAAAGGAGACTAGATTGCCTTTGTTTTGATGTTTGTGGCCTCAGAATT

GATCATTTTCCCCCCACTCTCCCCACACTAACCTGGGTTCCCTTTCCTTC

CATCCCTACCCCCTAAGAGCCATTTAGGGGCCACTTTTGACTAGGGATTC

AGGCTGCTTGGGATAAAGATGCAAGGACCAGGACTCCCTCCTCACCTCTG

GACTGGCTAGAGTCCTCACTCCCAGTCCAAATGTCCTCCAGAAGCCTCTG

GCTAGAGGCCAGCCCCACCCAGGAGGGAGGGGGCTATAGCTACAGGAAGC

ACCCCATGCCAAAGCTAGGGTGGCCCTTGCAGTTCAGCACCACCCTAGTC

CCTTCCCCTCCCTGGCTCCCATGACCATACTGAGGGACCAACTGGGCCCA

AGACAGATGCCCCAGAGCTGTTTATGGCCTCAGCTGCCTCACTTCCTACA

AGAGCAGCCTGTGGCATCTTTGCCTTGGGCTGCTCCTCATGGTGGGTTCA

GGGGACTCAGCCCTGAGGTGAAAGGGAGCTATCAGGAACAGCTATGGGAG

CCCCAGGGTCTTCCCTACCTCAGGCAGGAAGGGCAGGAAGGAGAGCCTGC

TGCATGGGGTGGGGTAGGGCTGACTAGAAGGGCCAGTCCTGCCTGGCCAG

GCAGATCTGTGCCCCATGCCTGTCCAGCCTGGGCAGCCAGGCTGCCAAGG

CCAGAGTGGCCTGGCCAGGAGCTCTTCAGGCCTCCCTCTCTCTTCTGCTC

CACCCTTGGCCTGTCTCATCCCCAGGGGTCCCAGCCACCCCGGGCTCTCT

GCTGTACATATTTGAGACTAGTTTTTATTCCTTGTGAAGATGATATACTA

TTTTTGTTAAGCGTGTCTGTATTTATGTGTGAGGAGCTGCTGGCTTGCAG

TGCGCGTGCACGTGGAGAGCTGGTGCCCGGAGATTGGACGGCCTGATGCT

CCCTCCCCTGCCCTGGTCCAGGGAAGCTGGCCGAGGGTCCTGGCTCCTGA

GGGGCATCTGCCCCTCCCCCAACCCCCACCCCACACTTGTTCCAGCTCTT

TGAAATAGTCTGTGTGAAGGTGAAAGTGCAGTTCAGTAATAAACTGTGTT

TACTCAGTGAA
