## Supplemental figure 2 for "Differential modulation of host cell Serine and Arginine-Rich (SR) protein and apoptotic factor BCL-2 gene expressions during Chandipura and Dengue virus infections"

Supplementary Figure 2 : SRp75 binding motifs in apoptotic factor BCL2 gene. Mtof sequences (TCCGGA) are highlighted in green color. Exons are in uppercase and introns are in lowercase letter.

CTCGATCCGGGCGATGGAGGAGGAAGCAAGCGAGGGGGCTGGTTCCTGAGCTTCGCAATTCCTGTGTCGCCTTCTGGGCTCCCAGCCTGCCGGGTCGCATGATCCCTCCGGCCGGAGCTGGTTTTTTTGCCAGCCACCGCGAGGCCGGCTGAGTTACCGGCATCCCCGCAGCCACCTCCTCTCCCGACCTGTGATACAAAAGATCTTCCGGGGGCTGCACCTGCCTGCCTTTGCCTAAGGCGGATTTGAATgtaggtggtgcgggggagcgggagtggggcgggggggactgcccagggagtgactttccgaggaaggcatttcggagaagacgggggta**gaaaa**ggctggtgggagattcagagtccactggtgctttcgatttgacttaagtgaagtatcttggaacctagacccagaccttcgtaagacccacaaagaaaccagttctggtacctggagggggaatggaatttttagggtaaatggcatgcatattaattatttttttttcctgaag CTCTTTCTCTCCCTTCAGAATCTTATCTTGGCTTTGGATCTTAGAAGAGAATCACTAACCAGAGACGAGACTCAGTGAGTGAGCAGGTGTTTTGGACAATGGACTGGTTGAGCCCATCCCTATTATAAAAATGTCTCAGAGCAACCGGGAGCTGGTGGTTGACTTTCTCTCCTACAAGCTTTCCCAGAAAGGATACAGCTGGAGTCAGTTTAGTGATGTGGAAGAGAACAGGACTGAGGCCCCAGAAGGGACTGAATCGGAGATGGAGACCCCCAGTGCCATCAATGGCAACCCATCCTGGCACCTGGCAGACAGCCCCGCGGTGAATGGAGCCACTGGCCACAGCAGCAGTTTGGATGCCCGGGAGGTGATCCCCATGGCAGCAGTAAAGCAAGCGCTGAGGGAGGCAGGCGACGAGTTTGAACTGCGGTACCGGCGGGCATTCAGTGACCTGACATCCCAGCTCCACATCACCCCAGGGACAGCATATCAGAGCTTTGAACAGGTAGTGAATGAACTCTTCCGGGATGGGGTAAACTGGGGTCGCATTGTGGCCTTTTTCTCCTTCGGCGGGGCACTGTGCGTGGAAAGCGTAGACAAGGAGATGCAGGTATTGGTGAGTCGGATCGCAGCTTGGATGGCCACTTACCTGAATGACCACCTAGAGCCTTGGATCCAGGAGAACGGCGGCTGGgtaagaaccaagccccttgtgtgtcccttttctttggcctctggtcagagatccccaacagcccttcttctgtatctctttctgttgtgttgggtgattgttggagacgttgatagttgaggaaacctgactggcctcatttcaccacaagaggttaactccttggaaatacaaatgaccctgtgcttcagagatcaggcatctttcaccctggtcacccccatgactccactattccatacttgtgttccagtttctcagcaaa**gaaaa**ctgtagcctatgtgttcattaagtcccaaacctgtgagggtaggaagttagttcggtgtttcccttaccttatacaatggttcacacacaaaggggtgatgagtataatgcctggagaagatgatgagcatcaatgacccacagacctctccatctgttccccccaaggggtctcagtcctatcctgggcttctctcccccaaatcaagttccctttatttcaaagtttgcgtgtatggcactctcctctcagactagggcatggccataggcaggagttcccactgggttgaatggctgagaattgtcagattttggggcagtgtaggctgtgcagattgtgtttgcactgcgccccctaagttcatttcccccaaagtgtgcgactttctgcatggttgccttgtttgtggagcacttgctgtgtcacacattaagccaggggactttgcacctcattgcctggggtcggttggggagctccagctgctctgcctgttactttccggccagagcccaggggcctcttcaagcactgtttgctggcaaagcaagaacctagtctttggggctctgtgggacattgagatttagtaacagacttcacatgcaggtataagttccagttctgtcatgtaactagctgtgtgaccttggacaggctgggttactttcttgaggcttagtttctttcactaaaaaatgtggcaagctctccttgccttacagtgtattgtaaatcttgggataatgtgagcacagtgactgacaccagcaaattccctttccagggagggagagttctcctgcccatcactgagccaggatgagatggtggtgaaagcttctttaccagtggtcactgtaggcaagttttgctcttcagtgaattcagtcctgttttcttgcttatctccctacctagcggaggtgcaggttttatgagagatgttctatcccctcagagtatgca**gaaaa**gcatcaagctgtg**gaaaa**taagtcatccattgaaggagctgaggtggaaggggacatgcaccatggactcaggaatgtcaagaatgcaactgtctataggctgggcctagggtgtggcaggcacttcactttaccactgagcaagagacctcctgagtttgccctggaaccccaccccctctctcattttgcatttctttgcaatttcaactcgtctgtgccagctcttgtaaaaggcaagggggtcaagtcaatagaggtcagataactggggtggggtcaggggaggtaaaggtggtgggttctcttgagctagctgactagacttgaaagccattctgtaaatgaactctctagggggataaatgattaaatagttcctgttggtttgtgcagtgacctctcctgttccctcctgctttgattctcttttggggcaaatatttgttcagctgattacaggagggacaggattccagtactgcagacacaccctctgaggcatctccaagtagggattgaggctgcttgcccactttgcctgactattgtctttggccagtggtcctaataccaggtgggttgttgccagctataaacaaaaggcagtctgggagaggtgcctggtctgctagccagcctctgcagaaggctacccctaccagagaattgccagctgcccagtggtgcatatgaaagaggtcttagacacagcaactctgggtaagtttattagatctccatggggttgaaattccatcttccatctctttgcttaagaatcatcagagcctgttggacaacataccacatgtcacttcctcttggtgaacagaggggatgtgtgtggaagagccagttatatcagaaccatttctgagcacacattctctccctaaggccagggagtaaagaacatttgactcaggatcctttgagcctgggtagaaggtacccgcccccttcctctcaggtattatcctgttccaagctgcaccgactggctactgtggtggcatcgggtaattatcctggagactgcggctgggagatccttaagggtaggagttatgggctattctactcagggctatgttctaaatttagcatccctacaggaaaggggaaggaaattagtctttattgagtacctgttaagtactaggcactgtgttaggtcctctacttatgcaaccagtgtttcttgagcatctactgtgaagctagcagtgtactaggtgctggccataagccacgcatggtaccttccttcatggaaacggcatgctgtttcatatattctacagtaaccttccaaggaaggtgagtttattcccagtttacacatgaaggtctttgtgcctaagtgactgaccccaggtgtccctcagccagttaagtgtctgaatgagaatttacaccaaggttggcataattctaaagccaacacttttcatcctatcaagttgtccagggagtaataaagtgactagacatgctggcccagaaagccagagatgctgttgggccaagcaaaccatgcctgtgaaggtgctcactctggactgataactggctttgttcaggtagctatgggaaggatagttggcagtagcaggttcccagaggcgtgttggtagcctcagattctggcactggtaaaactcagacatggcaggtttcaagtgggcatcagggttttgcccagagtgagctgcttattaattttttctttttttttttttttttgagacggagtctcgctcttgttgcctaggctggagtgcagtggcgcgatctcagctcactgcagcctctgcctcccgggttcaagtgattctcctgcctcagcctcccaagtagctgggattacaggcatgcgccaccacacctggctaattttgtatttttagtagagatggggtttctccatgttggtcaagctggtttcaaactcccaacctcaggtgatccgcctgcctcagcctcccaaaatgctgggattacaggcatgaaccaccgtgcctggccccggccaattttttcattgttaaaaccctctcaggccctgccaatagatgacagtggtgtgcgtgtggggagaaggtggcctgccccaggtatctctacttagtggtggagtgatgcttcaaggactgaaagtgtagacccttagccagattactgcctgcctggagtcacctgatgaaatgctgaacctgttcggctgtcttgtccatctgtctctcccgcctgtaaaatggaataatgatagccagctgccatagcttgactcgagtaaaggatctagcacaaaagcccttctcctgccataattgtagggtggaaaccctttaatgaatgggcagcttccctcagagaactgactcttccccctagagtttctccttgagaaacaaagtccctgtgatactttcctggaatgttgtatacatgaccttccccgaagggacacaagtgtttctggtgctttccaatgggaatgtgggaagggacccaggtgggccttgccactttgggattgctgtccctgaagaaatcccttagcctgatagaaacgtaattgttgggagcaatgaactgtgttgggggagaaaacataacttggcctttcttaagctgtatggctcagtggtctgagtttctgtagatctcttattgcctaatgtggcttcttcagatggatgtatgcttagttgaagcacctcagctccctttgcagatagtggcagttctgtggttttcttagggaaaccccattagaaaccccaaatgtggaaatttttagtaatctcattcgggtcccaaaattaagtagaagtgtctcaggggctcccccttgaaacacctacttcttctgagtgtacatttgaggttgctcttaaaattctgttaccatcacagggttttccagacttcctaggaataatttccccagtagccccacagagccctttagcttctgtcgaaaattacccgggtattttttcagggggctccaatggaacctggagggttttcttgacttttgtatgggtgtccccagatttgtttgggctgcatgaaacagcagttgttatctctgtcccttcccccagcagatatctgtcctgaggccagtgggcttgagcgtcctcctctgtgtccagtctccctggttttgccagtgccctcagccttccatcatggggctttcttgaagcaactgtggttttgcctgagtttaatgtggatagacataggggtaattcccagtgtactgcttttctacggaagtataatctcctactgccttaattaagcagaggccctgtccccacccaacctccaacctaattccacagagtcctccctaaggactccctcttttttctttgaaggagtcactctactcctggatcctcttctcttctgcccctctatccgacagtctgtcttccttgttcccagttggcagacctgagtcaggcaatcccactgggtctttttgtttagcctggggaagagaagacccaggagaatatcaccagcttcagatatctgaaggaagagggactgagtctttttttattggtgtggtacctggcaaaacttggataaatgaaaagaagttaacagttggtagagcttaaattatgtgcagagaagacatttctaacagatttactctgtcctagatttaggagctcctcagggtggaggccacatcttgtccatgcttgtttcctcagcactgagcagaggccttgggcatagcagatacctaattgaagagaaacccactaaatttctgaagaaacagacccttgtcaagccagaataacaataatgggagtggctatttacctaataattcattcgttgagtcaaacaatgtttattgagctcctactgtgtaccatactgggagtatggcagtaaacaaaagtaagcaaaactaagtttctgtcctcatggagctcacattccaatgaggagtgacaggcaataaatatataattacatgtttatatcagatgattgtacatgctatgaagaaaataaagcaggttggggcatagggcatgtctagcaagggagggtggggctgcagttttaaaagagtgatcaaagaaggtgtaagggagaaaaaaatggctttcctctacccttctaggtcctttggctgggctgagaattaaattgacatagacagattaactggaggaaaaataattaattacatagttgtacacaggaatcctacaaaatatgaggctcaaagaagggtcagataattgaagcttatatagcatcctgagctacagaaaagaatatggggcttgggacttctgttaggggcttgagtggcagcaacccaagttatggcagggtgagggaaagaattggtggtggcgtgtgtctgtggtcccagatactgttgagggggtgtactgaggtgggaggatcacttgagcctgggaggtggaggttgcagtgagccgagatcaagccactgcgctccaacctgggtgacagagcaagaccctgtctcaaaaaataaaaataaaaagtctctctggtaataaaatttgtctggagcagccctcttcctaatacagatacctttactagtgtagatttcctttttagatgtgaatttcttttacaaaaggacagcttttcagagctactcttatgtctgcagtttctcagaataaccagttcaaaatatgccaaagaagtatatttttggggtggcatattgttttcctacagtcatattttgggatggtatgtcctgagtcccaacaaaagtattacctaaatttatttgagccaagcgctaaatgaggtaaggaggtggatgatgtaggtaccctacagcaaagggtgtttaaagcagagggaatggactcaggagggagcatgcttgacattttcaaggatgagcaagaaggtcaatgtggctggagaagaaacagcaaaggggaaaagggtagaaggtggggagggaatggttgaggaggaggatggagaagtaggtgggggccagatcgcatacagcattacagaccatgatgaggatttggctttggtggaatggtctgtattccttttttttttttttttgag

acagagtctttctctgtctcccagactggagtgcagtggcacgatctctg

TACTCAGTGAA
